# Chimeric a-subunit isoforms generate functional yeast V-ATPases with altered regulatory properties *in vitro* and *in vivo*

**DOI:** 10.1101/2022.02.04.479146

**Authors:** Farzana Tuli, Patricia M. Kane

## Abstract

V-ATPases are highly regulated, multi-subunit proton pumps that acidify organelles. The V-ATPase a-subunit is a two-domain protein containing a C-terminal transmembrane domain that participates in proton transport and a N-terminal cytosolic domain (aNT) that acts as a regulatory hub, integrating environmental inputs to regulate assembly, localization, and V-ATPase activity. Tissue- and organelle-specific a-subunit isoforms exist in most organisms, but how regulatory inputs are decoded by aNT isoforms is unknown. The yeast *S. cerevisiae* encodes only two organelle-specific a-isoforms, Stv1 in the Golgi and Vph1 in the vacuole. Based on recent structures, we designed chimeric yeast aNTs in which the globular proximal and distal ends are exchanged. The Vph1 proximal-Stv1 distal (VPSD) aNT chimera binds to the glucose-responsive RAVE assembly factor *in vitro* but exhibits little binding to phosphoinositide lipids that activate V-ATPases. The Stv1 proximal-Vph1 distal (SPVD) aNT lacks RAVE binding but binds more tightly to phosphoinositides than Vph1 or Stv1. When attached to the Vph1 C-terminal domain *in vivo*, both chimeras complement growth defects of a *vph1*Δ mutant, but only the SPVD chimera exhibits wild-type V-ATPase activity. Cells containing the SPVD chimera adapt more slowly to a poor carbon source than wild-type cells but grow more rapidly than wild-type after a shift to alkaline pH. This is the first example of a “redesigned” V-ATPase with altered regulatory properties and adaptation to specific stresses.

## INTRODUCTION

The intracellular compartments of eukaryotic cells maintain a unique pH range which is critical for their function (Casey et al., 2010). The organelles of the secretory and endocytic pathways become progressively more acidic, with the lysosome/vacuole as the most acidic organelle (Huotari & Helenius, 2011). All acidic compartments in these pathways possess Vacuolar H^+^-ATPases (V-ATPases) which acidify their lumen. In addition, some specialized cells have plasma membrane V-ATPases that secrete H^+^ into the extracellular space. V-ATPases play important roles in many cellular processes including vesicular transport, neurotransmitter loading, ion homeostasis, urine acidification, and bone resorption (Breton & Brown, 2013; Collins et al., 2020; Eaton et al., 2021; Forgac, 2007). Consistent with their diverse roles in cell physiology, aberrant activity of V-ATPases has been associated with many pathological conditions such as osteopetrosis (Frattini et al., 2000), renal tubular acidosis (Smith et al., 2000) and metastatic cancer (Sennoune et al., 2004). However, complete loss of V-ATPase activity is lethal in higher eukaryotes (Sun-Wada et al., 2003), which makes therapeutic targeting of V-ATPases very difficult (Kane, 2012).

V-ATPases are ATP hydrolysis-driven proton pumps that are conserved from yeast to humans. These multi-subunit protein complexes consist of two main subcomplexes where the cytosolic, peripheral complex, V_1_, hydrolyzes ATP and membrane-bound complex, V_o_, contains the proton pore (Forgac, 2007). V-ATPases undergo different modes of regulation to fine tune their activity depending on cellular need (Forgac, 2007). One major mechanism of regulation of activity is by reversible disassembly where the V_1_ and V_o_ sectors reversibly disassemble in response to many signals such as upon glucose deprivation (Kane, 1995) (Sumner et al., 1995). Signal-dependent disassembly of V_1_ from V_o_ leads to the inactivation of both ATP hydrolysis (Parra et al., 2000) and proton pumping (Couoh-Cardel et al., 2015). Significant conformational changes of the disassembled V_1_ and V_o_ complexes prevent reassembly when enzyme activity is not needed. The RAVE (Regulator of acidification in vacuole and endosome) complex is required for rapid reassembly of yeast V-ATPase *in vivo* (Seol et al., 2001). V-ATPase activity can also be modulated by phosphatidylinositol phosphate (PIP) lipids (Banerjee et al., 2019; Vasanthakumar et al., 2019) which are enriched in organelle membranes and may stabilize the assembled conformation of V-ATPase. In addition, isoforms of many V-ATPase subunits allow for tissue-specific and organelle-specific regulation. How isoform composition endows V-ATPases with distinct catalytic and regulatory properties is poorly understood.

The a-subunit from the V_o_ sector is encoded by multiple tissue- and organelle-specific isoforms in most eukaryotic cells. In yeast, the only V-ATPase subunit isoforms are in the V_o_ a-subunit and are encoded by the *VPH1* and *STV1* genes. Vph1-containing V-ATPases are enriched in the vacuole (Manolson et al., 1992), and Stv1-containing V-ATPases localize to the Golgi (Manolson et al., 1994). These isoforms determine localization of their respective complexes and confer different catalytic and regulatory properties. In all eukaryotes, a-subunit isoforms consist of a cytosolic N-terminal domain (NT) and a membrane bound C-terminal domain (CT). Chimeras of yeast a-subunits containing NT and CT of the opposite isoforms revealed that the NT domain encodes information for localization and reversible disassembly (Kawasaki-Nishi et al., 2001). Subsequent studies showed that the NT domains also dictate binding to different organelle-enriched phosphoinositide lipids (Banerjee et al., 2019; Banerjee & Kane, 2017; Li et al., 2014). The overall backbone structure of the NT domain of eukaryotic a-subunit isoforms is predicted to be similar, with 50-70% similarity at the amino acid level. Cryo-EM structures reveal that the aNT domains are characterized by a proximal end (which connects to the aCT domain) and a distal end; the two ends are connected by a long coiled-coil (Figure 1). Recent structures of Vph1-containing V_o_ and Stv1-containing V_o_ revealed strong similarity in overall structure but failed to resolve poorly conserved loops at both proximal and distal ends of aNTs (Vasanthakumar et al., 2019). These loops are likely flexible and may contribute to unique catalytic and regulatory properties of the isoforms. Indeed, sequences from these loop regions have previously been implicated in lipid binding (Banerjee et al., 2019; Banerjee & Kane, 2017).

**Figure 1:**
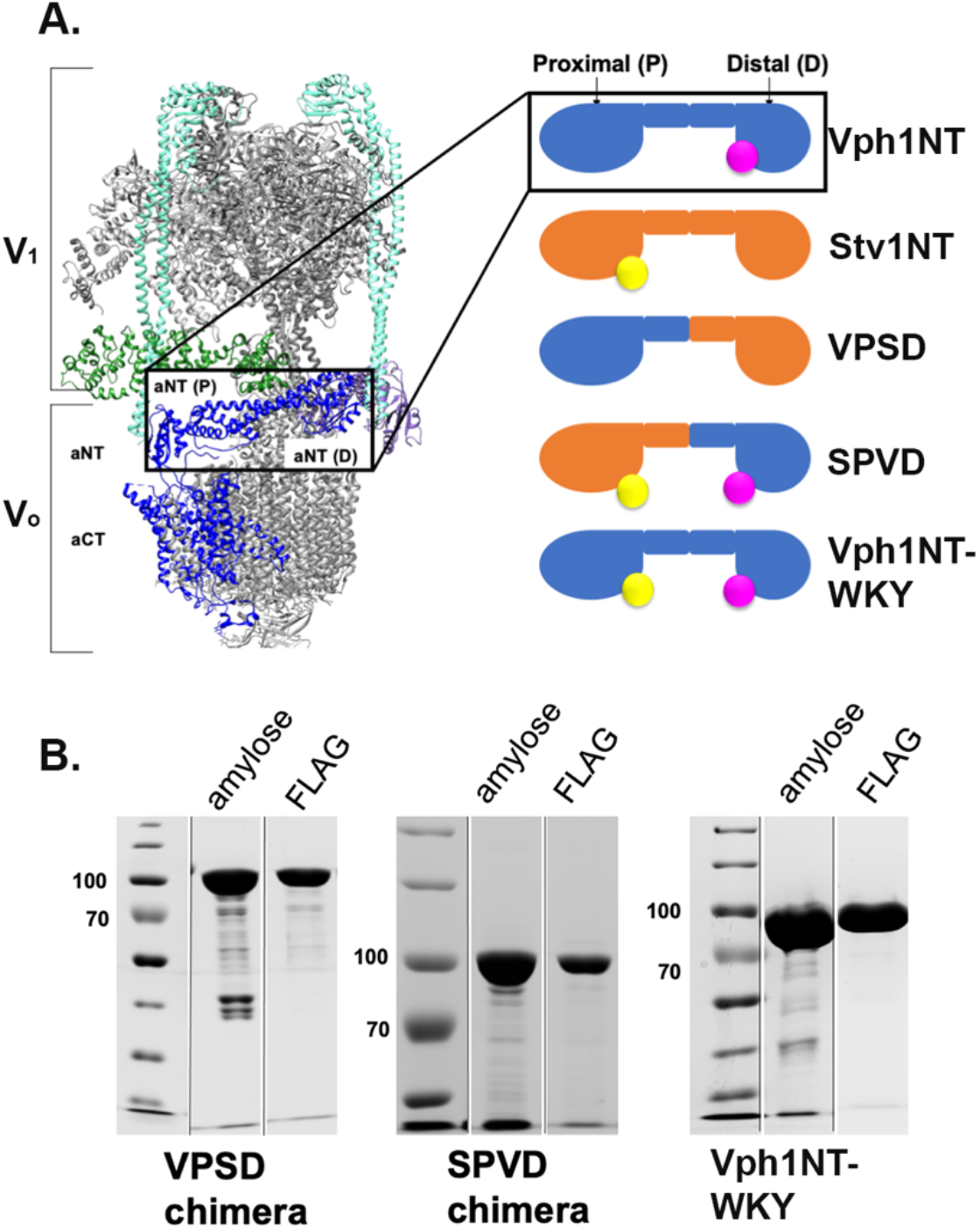
Design and construction of chimeric and mutant a-NT’s. **A. Left**, Cryo-EM structure (3.1-Å resolution) of human V-ATPase holoenzyme (PDB 6WM2) where the a-subunit is shown as a dark blue polypeptide chain. Subunits that interact with the aNT domain are represented in color; V_1_ E and G subunits are in turquoise, V_1_ H-green, V_1_ C-purple. The aNT domain is boxed, and the proximal (P) and distal (D) ends are indicated. **Right**, Schematic diagrams of yeast Vph1NT and Stv1NT with the proximal and distal ends indicated along with chimeric proteins constructed by swapping the two ends between the isoforms. The Vph1NT-WKY mutant has only a 6 amino acid sequence from Stv1NT added to the proximal domain. Amino acids previously implicated in PI(3,5)P_2_ activation of Vph1-containing ATPases are indicated in magenta, and the amino acids implicated in PI(4)P binding to Stv1 are shown in yellow. **B**. Coomassie stained SDS–PAGE showing MBP-tagged VPSD and SPVD chimeras and Vph1NT-WKY mutant after the first (amylose) and second (anti-FLAG) affinity purification steps. Lines indicate positions where non-adjacent lanes from the same gel were moved together.

The aNT domain is an important regulatory hub of V-ATPase for several reasons. First, it occupies a critical position at the V_1_-V_o_ interface **(Figure 1A)**. It undergoes significant conformational changes in the V-ATPase holoenzyme vs. disassembled V_o_ subcomplexes (Couoh-Cardel et al., 2015). Second, the proximal and distal ends form quaternary interactions with different V_1_ subunits (Oot & Wilkens, 2012; Sharma et al., 2018); these interactions must be supported by all isoforms to preserve V-ATPase assembly and activity. Third, the aNT harbors many different regulatory interactions. Cellular factors important for V-ATPase regulation such as the RAVE complex (Smardon et al., 2014), PIP lipids (Banerjee et al., 2019; Banerjee & Kane, 2017; Li et al., 2014), glycolytic enzymes (Lu et al., 2007; Su et al., 2008), and the Rab-GEF protein ARNO (Hurtado-Lorenzo et al., 2006) all interact with aNT domains. However, it is not well understood where the information for these regulatory interactions resides in the aNT domains.

Here, we aimed to understand where two of the regulatory interactions, RAVE and PIP lipids, reside in the aNT domain and how the information from these two inputs is integrated to control V-ATPase activity. Toward this goal, we have constructed chimeric aNTs by swapping the proximal and distal ends between Vph1 and Stv1. *In vitro* studies with the resulting chimeras suggest that the information for RAVE binding resides in the proximal end of Vph1NT. Lipid binding experiments suggest that multiple sequences in Stv1NT from both ends promote PI(4)P binding. Unlike previous chimeric a-subunits, a chimera containing proximal end of Stv1 in combination with distal end of Vph1 assembles into fully functional V-ATPases in vacuoles when attached to the Vph1 C-terminal domain. However, the altered regulatory properties of this chimera have significant physiological consequences for yeast cells.

## RESULTS

### Design and expression of chimeric and mutant proteins

In order to better understand the regulatory information embedded in the cytosolic aNT domains, we generated chimeras of Vph1NT and Stv1NT *in vitro* by swapping the proximal and distal ends between the two isoforms **(Figure 1, Supplemental Figure 1A)**. Pairwise alignment of the two isoforms helped define domain boundaries for chimera generation (**Supplemental Figure 1B**). We placed the chimeric junction at a region of high sequence similarity in the middle of the coiled-coil. Medium resolution cryo-EM structures of intact V-ATPases containing both Vph1 (Benlekbir et al., 2012; Zhao et al., 2015) and Stv1 (Vasanthakumar et al., 2019) are available, but there are regions of the aNT domains that are missing from the structures or poorly resolved, including the regions previously implicated in lipid binding (Banerjee et al., 2019; Banerjee & Kane, 2017). High confidence models for Vph1NT and Stv1NT (**Supplemental Figure 1A)** suggest that the design of the chimeras should preserve the structures of the proximal and distal ends of each isoform. Previous work identified a binding site for PI4P in the proximal domain of Stv1NT that is indicated by a yellow circle (Banerjee & Kane, 2017), and binding site for P(3,5)P2 in the distal domain of Vph1NT that is indicated by a pink circle (Banerjee et al., 2019) in the **Figure 1A** and **Supplemental Figure 1A**. The resulting chimeras were designated Vph1NT_proximal_ Stv1NT_distal_ (VPSD) and Stv1NT_proximal_ Vph1NT_distal_ (SPVD) **(Figure 1A)**. In addition to these chimeras, we constructed a Vph1NT-WKY mutant in which we added the Stv1NT <u>W83KY</u>ILH sequence containing the previously identified (underlined) PI(4)P recognition site (Banerjee & Kane, 2017; Finnigan et al., 2012) into the corresponding loop of the Vph1NT proximal domain.

The chimeric and mutant aNT proteins were expressed in E. coli with a N-terminal maltose binding protein (MBP) tag and a C-terminal FLAG tag and purified through sequential amylose and anti-FLAG affinity chromatography **(Figure 1B)** followed by gel filtration. The gel filtration profiles of the chimeric and mutant proteins **(Supplemental Figure 1C)** were very similar to those of the wild-type aNTs, including the characteristic monomer-dimer peak that is observed for wild-type Vph1NT and Stv1NT (Banerjee & Kane, 2017; Oot & Wilkens, 2012). The monomer fractions were used in the subsequent assays.

### The proximal end of Vph1NT binds to Rav1 (679-898)

The yeast RAVE complex is an isoform-specific assembly factor required only for the assembly of Vph1-containing V-ATPases; Stv1-containing V-ATPases can assemble independently of RAVE (Smardon et al., 2014). Vph1NT directly interacts with one RAVE subunit, Rav1, in two-hybrid and pull-down assays (Smardon et al., 2014; Smardon et al., 2015) (Jaskolka & Kane, 2020). A central fragment of Rav1 (aa 679-898) has a binding site for Vph1NT (Smardon et al., 2015) and point mutations and small deletions in this region abolish recruitment of RAVE to the vacuolar membrane *in vivo* (Jaskolka & Kane, 2020). Expressed Vph1NT co-elutes with a Rav1(679-898)His_6_ fragment from TALON resin (Jaskolka & Kane, 2020; **Figure 2A;**). In contrast, no coelution of Stv1NT with Rav1(679-898)His_6_ was observed **(Figure 2B)**, consistent with pull-down results using an overlapping fragment (Smardon et al., 2014). Although these experiments support isoform-specific binding, chimeric aNTs provide an opportunity to narrow down the RAVE binding domain. We performed similar pull-down assays with the Rav1(679-898)His_6_ fragment and the chimeric aNTs. We found that the VPSD chimera coeluted with Rav1(679-898) His_6_ **(Figure 2C)**. In contrast, the SPVD chimera, which lacks the proximal end of Vph1NT, did not coelute with the Rav1(679-898) His_6_ fragment **(Figure 2D)**. (Neither chimera was detected in controls without the Rav1 fragment, indicating the interaction is specific.) These data suggest that the Vph1 proximal end present in the VPSD chimera has the information for binding to this critical region of Rav1, whereas the distal end is not important for the interaction. As indicated in the scheme in Figure 2C, this places Rav1 binding (indicated by a star) at the opposite end from the PI(3,5)P2 binding region.

**Figure 2:**
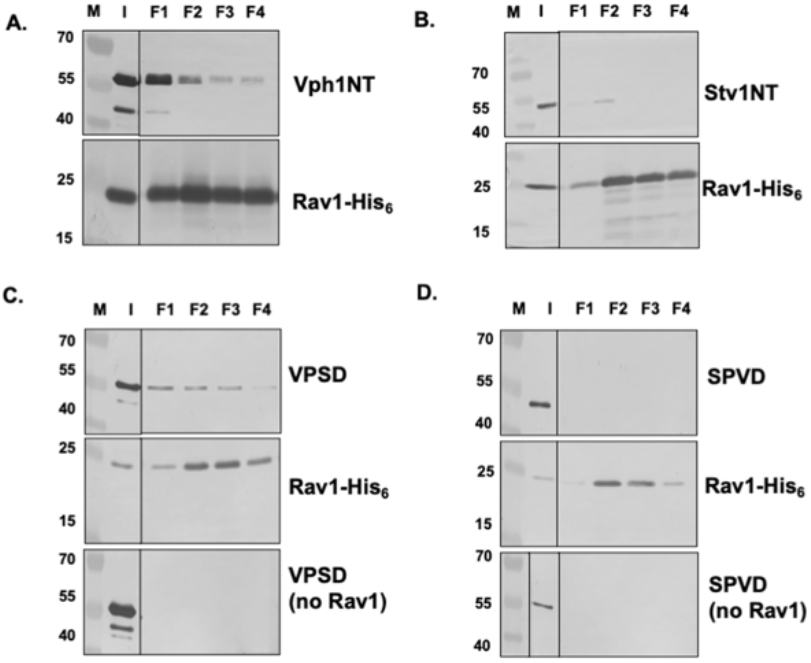
The proximal end of Vph1NT binds to Rav1 (679-898). Vph1NT, Stv1NT, and the VPSD and SPVD chimeras were expressed and purified as in Figure 1 except that the MBP tag was removed with Prescission protease. Rav1 (679-898)His_6_ was expressed and purified as described (Smardon et al., 2014). All four proteins were brought to a concentration of ∼1 μM, then were mixed with excess Rav1 (679-898)His_6_ at a 5:1 molar ratio. The mixtures were added to TALON resin and incubated at 4°C for 2 h. to allow binding. The pull-down assay was performed as described (Jaskolka & Kane, 2020). The Rav1 (679-898)His_6_ fragment was used to pull-down: **A**. Vph1NT, **B**. Stv1NT, **C**. VPSD and **D**. SPVD, and the bound proteins eluted with imidazole in the indicated fractions. No Rav1 control, for each pull-down assay the indicated protein was also mixed with TALON resin in absence of Rav1, data not shown for Vph1NT and Stv1NT. Anti-FLAG monoclonal antibody was used to probe for Vph1NT, Stv1NT, VPSD and SPVD while anti-His_6_ antibody was used to probe Rav1 (679-898)His_6_. Lines indicate positions where non-adjacent lanes from the same gel were moved together. M: molecular mass markers, I: input.

### Transfer of PIP specificity between aNT isoforms

PIP lipids interact with V-ATPase isoforms and contribute to their activity, localization, and regulation (Banerjee et al., 2019; Banerjee & Kane, 2017; Li et al., 2014). We tested binding of the chimeric yeast aNT constructs to PI(4)P and PI(3,5)P_2_ in a quantitative liposome pelleting assay (Chandra et al., 2019). Liposomes with or without added phosphoinositide were incubated with a defined concentration of the aNT protein. Liposomes and bound protein were isolated by centrifugation as described in Materials and Methods, and binding was assessed by measuring the proportion of total protein in the pellet. PIP-specific binding was determined by subtracting the protein bound in the absence of phosphoinositide from that bound in the presence of 5% phosphoinositide. All of the proteins were tested for pelleting in the absence of lipid, and none pelleted, indicating that the presence of protein in the pellet was liposome-dependent.

Stv1NT binds to Golgi-enriched PI(4)P *in vivo* and to PI(4)P-containing liposomes in a liposome flotation assay (Banerjee & Kane, 2017). Mutation of K84, located in the proximal end of Stv1NT, abolished binding to PI(4)P-containing liposomes *in vitro*, suggesting this region is necessary for lipid binding. It was not determined whether this region is sufficient for PI(4)P interaction, and the contribution of other regions of Stv1NT to lipid binding is not known. As shown in **Figure 3A**, Stv1NT showed much more binding to PI(4)P liposomes than Vph1NT at a protein concentration of 0.5 μM. We tested whether we could transfer Stv1NT PI(4)P specificity to Vph1NT using the Vph1NT-WKY mutant (**Figure 1 schematic**). As shown in **Figure 3A**, this mutant bound to PI(4)P liposomes comparably to Stv1NT and significantly better than Vph1NT. However, the SPVD chimera, in which the entire proximal end domain of Vph1NT was replaced with Stv1NT, bound even more strongly to PI(4)P suggesting that other sequences in the Stv1NT proximal domain may strongly enhance binding **(Figure 3A, B)**. The VPSD chimera, in which the entire distal end domain of Vph1NT was replaced with the distal end of Stv1NT also showed significantly better binding than Vph1NT alone at 0.5 μM protein **(Figure 3A, B, C)**. Together these data support the previously identified WKY sequence as a major PI(4)P binding site in Stv1 which is sufficient to transfer PI(4)P binding to Vph1NT. However, both the remainder of the proximal and the distal ends of Stv1NT can further promote PI(4)P binding.

**Figure 3:**
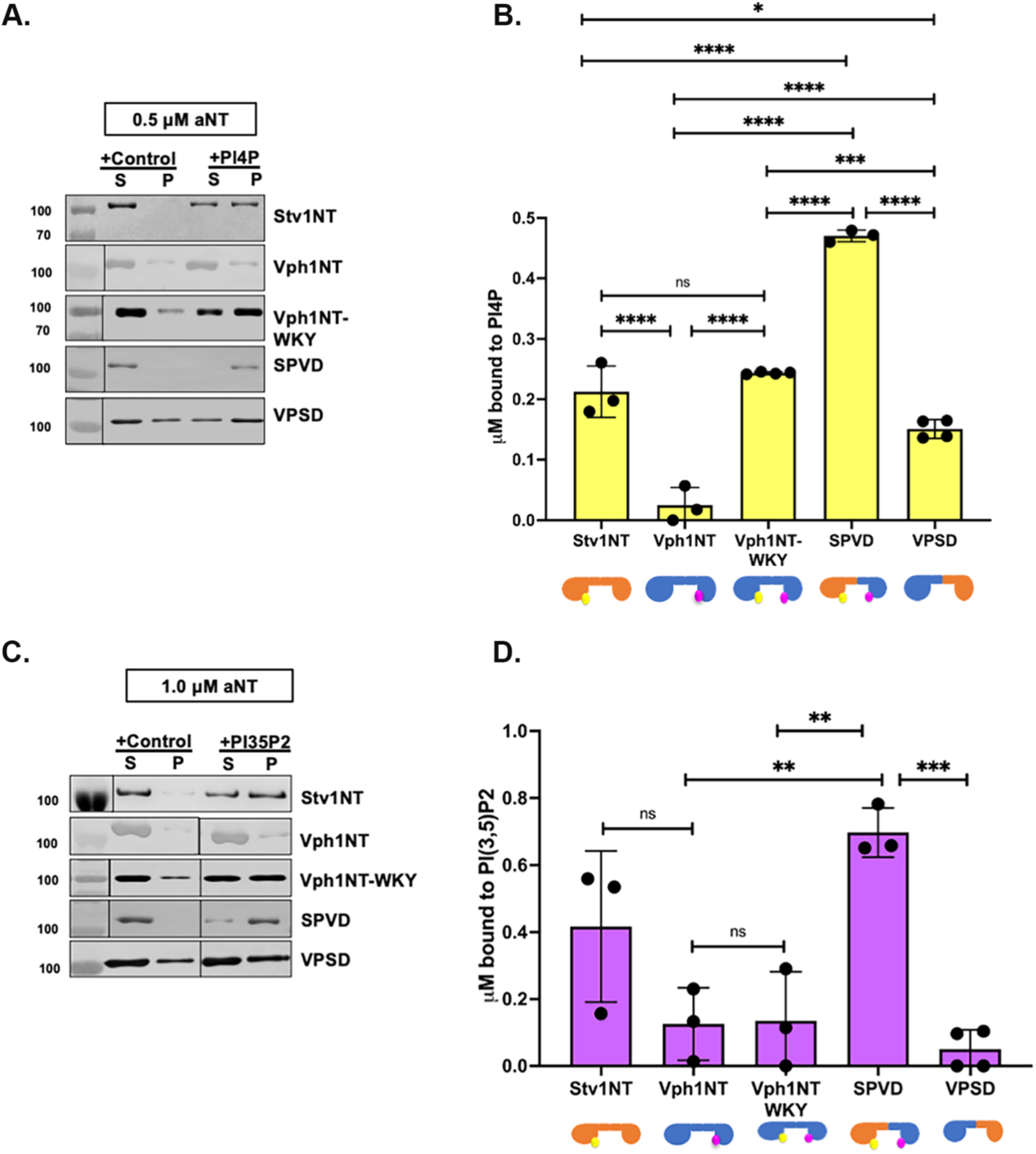
PIP lipid binding of chimeric and mutant aNTs in liposome pelleting assays. Liposomes were prepared as described (Banerjee & Kane, 2017). All proteins were expressed and purified as in Figure 1. Protein solutions at a final concentration of either 0.5 μM or 1 μM (as indicated) were incubated with control (no PIP), 5% PI(4)P or 5% PI(3,5)P_2_ containing liposomes (final concentration of 0.33 mM) in separate centrifuge tubes as described in *Materials and Methods*. Following centrifugation, the supernatant (S) and pellet (P) fractions were collected and resuspended in 100 μl of 50 mM Tris-Cl, 100 mM NaCl (pH 7.4) before TCA precipitation and analysis by SDS-PAGE and immunoblotting with anti-FLAG antibody. **A**., **C**. Representative immunoblots showing distribution of the indicated proteins between the supernatant (S) and pellet (P) fractions from samples containing control and PIP-containing liposomes. **A**. 0.5μM of each protein with control and PI(4)P-containing liposomes, **C**. 1 μM of each protein with control and PI(3,5)P_2_ containing liposomes. In **A** and **C**., vertical lines represent movement of non-adjacent lanes from the same gel next to each other. **B, D** At least three assays were performed for each protein-liposome combination and band intensities were determined using ImageJ. PIP-specific binding was determined by subtracting the proportion of total protein bound in the control pellets from the proportion in the PIP pellet, then multiplying by the total protein concentration to get PIP-specific binding. Bar graphs show the average concentration of each protein bound specifically to liposomes containing the indicated PIP lipid. **B**. PI(4)P-specific binding at total protein concentrations of 0.5 μM **D**. PI(3,5)P_2_-specific binding at total protein concentrations of 1 μM. Bar graphs represent mean ± SD; each dot is a biological replicate. In each graph, binding for each protein was compared to every other protein by ordinary one-way ANOVA with Tukey’s multiple comparison test; significant differences between the strains are indicated: ****, P<0.0001 ***, P<0.0005; **, P<0.005; *, P<0.05; ns, P>0.05

Direct *in vitro* interaction between the vacuolar isoform Vph1NT and PI(3,5)P_2-_containing liposomes has not been observed previously although Vph1NT shows PI(3,5)P_2_- dependent recruitment to membranes *in vivo* (Li et al., 2014). Furthermore, addition of exogenous PI(3,5)P_2_ specifically enhances the ATPase and proton pumping activities of Vph1-containing V-ATPases in isolated vacuolar vesicles and this enhancement is abolished by mutations in the Vph1 distal domain (Banerjee et al., 2019). We found little binding of Vph1NT or the chimeric proteins to PI(3,5)P_2_-containing liposomes in a liposome pelleting assay at 0.5 μM protein (the concentration used in Figure 3A), so we conducted the assay in the presence of 1 μM protein **(Figure 3B)**. Under these conditions, the SPVD chimera showed significantly higher binding than Vph1NT, the Vph1NT-WKY or VPSD chimera.

### Expression of chimeric aNTs as part of intact V-ATPases

Although the expressed aNT domains are optimal for *in vitro* assays, they cannot capture the full function of the aNT domains *in vivo*, where they are tethered in proximity to membranes by attachment to the integral membrane aCT domains and must bind to multiple V-ATPase subunits. To compare their functions as part of the intact enzyme, chimeric aNT sequences were integrated into *VPH1*, replacing the Vph1NT sequence and placing them adjacent to the Vph1CT sequence and under control of *VPH1* promoter as depicted in **Figure 4A**. The resulting strain still has wild-type Stv1 localized to the Golgi and would be expected to express the chimeric a-subunits at higher levels since the *VPH1* promoter is much stronger (Kawasaki-Nishi et al., 2001; Manolson et al., 1994). We performed growth assays to assess V-ATPase function in the yeast strains containing chimeric a-subunits. Mutants lacking V-ATPase activity grow best at pH 5. Loss of Vph1 function makes yeast sensitive to high extracellular pH and elevated level Zn^2+^ concentrations (Manolson et al., 1994) as shown for the *vph1*Δ mutant. However, both chimeric aNT mutations complement the growth defects of the *vph1*Δ mutant, suggesting they are tolerated *in vivo* **(Figure 4B)**.

**Figure 4:**
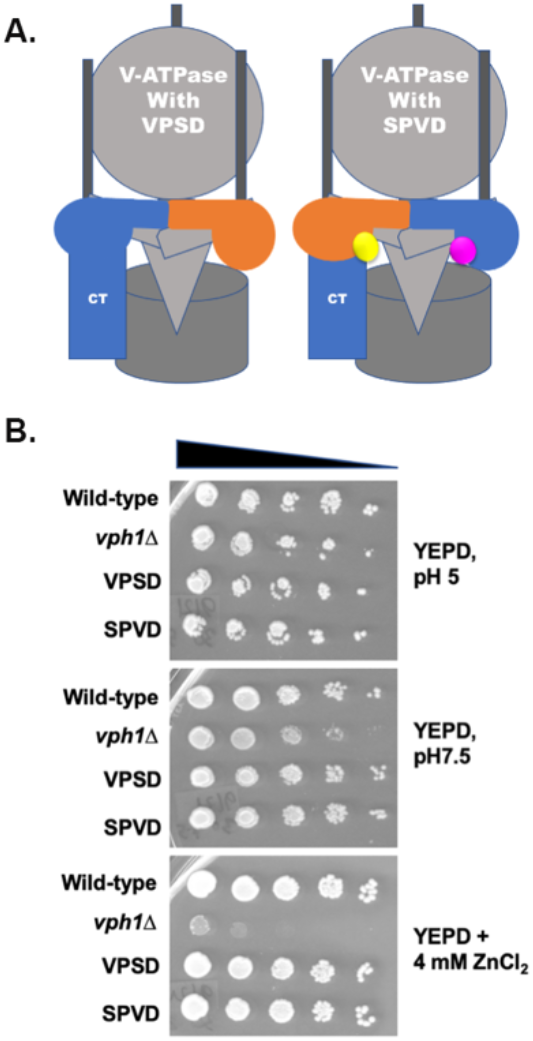
Chimeric aNT domains function as part of the V-ATPase complex. **A**. Diagrams depicting intact V-ATPases containing VPSD or SPVD chimeras replacing Vph1NT as part of the intact a-subunit. PIP binding sites are depicted as in Figure 1. **B**. Growth of wild-type, VPSD and SPVD strains was compared to that of a *vph1*Δ strain by growing each strain to log phase in YEPD pH 5, diluting to an A_600_ of 0.8 then serially diluting 10-fold in a microtiter plate. Cells were transferred to YEPD pH 5, YEPD pH 7.5 and YEPD Zn^2+^ plates and grown for 3 days at 30°C.

### The V-ATPases containing SPVD chimera are fully assembled and active

We isolated vacuolar vesicles from wild-type cells and the strains containing the SPVD and VPSD chimeras and measured V-ATPase activity and assembly at the vacuole. Vacuolar vesicles from the VPSD chimera had significantly less V-ATPase activity than the wild-type or SPVD vesicles (**Figure 5A)**. In order to better understand the source of the defect in the VPSD vesicles, we compared the levels of the V_1_ subunits A, B, and C, V_o_ subunit d, and control vacuolar membrane protein ALP between the vesicle preparations. Although alkaline phosphatase (ALP) levels are similar between the vesicle preparations, indicating comparable purity, all of the V_1_ subunits and V_o_ subunit d are at lower levels in the VPSD vesicles, suggesting a V-ATPase assembly or stability defect **(Figure 5B)**. The chimeric a-subunits showed different interactions with antibodies against Vph1 and Stv1. One of the monoclonal antibodies (7B1H1) raised against yeast vacuolar membranes (Kane et al., 1992) not only recognized Vph1 as expected, but also cross-reacted with the SPVD chimera. In contrast, another monoclonal antibody (10D7) only recognized Vph1 and VPSD, but not SPVD. A polyclonal antibody against Stv1 recognized VPSD and SPVD but not Vph1.

**Figure 5:**
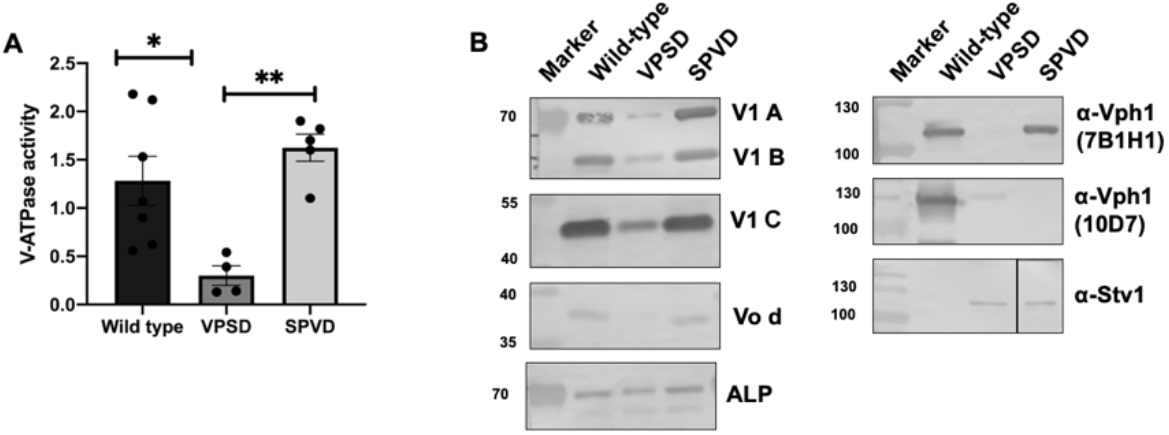
The V-ATPases containing SPVD chimera are active and fully assembled: Vacuolar vesicles were isolated from wild-type cells and cells containing the VPSD and SPVD chimera. **A**. V-ATPase–specific activity (defined as the activity sensitive to 100 nM concanamycin A and expressed as μmol/min/mg protein) was measured for at least four independent preparations of vacuolar vesicles, and the mean ± SEM is shown. **B**. The vesicles were solubilized, separated by SDS-PAGE, transferred to nitrocellulose, and probed with antibodies against V_1_ subunits A, B, and C, V_o_ subunit d, or alkaline phosphatase (ALP), a vacuolar integral membrane protease. Vesicles were also probed with two monoclonal antibodies specific for Vph1 (7B1H1 and 10D7) as well as polyclonal antisera against Stv1; none of these antibodies could detect the a-subunit from all three strains. For the detection of A- and B-subunit 2.5 μg of vacuolar protein was loaded. 10 μg of protein was loaded to detect the C subunit, and 7.5 μg of protein was loaded for visualization of a-subunits and alkaline phosphatase.

In contrast to the VPSD chimera, vesicles from the SPVD chimera had a mean ATPase activity slightly higher than those from wild-type cells, although the difference in activity was not statistically significant. The levels of V_1_ subunits in the SPVD vacuolar vesicles were also as high, or slightly higher, as in wild-type vesicles, and the V_o_ d-subunit was at similar levels. We also measured proton pumping in the wild-type and SPVD vesicles and observed that coupling ratios (the ratio of the initial rate of proton pumping to the initial rate of ATP hydrolysis) were also very similar between wild-type and SPVD vesicles (data not shown). In summary, despite differences in regulatory properties, the SPVD mutant is as assembled and active as the wild-type V-ATPase. These data indicate that, for the first time, we have generated a fully functional V-ATPase predicted to have significantly different regulatory properties than wild-type. We proceeded to test the physiological impact of these regulatory differences.

### The SPVD chimera is partially independent of RAVE function

The RAVE complex plays roles in both biosynthetic assembly of the V-ATPase and reassembly after disassembly induced by glucose deprivation. *rav1*Δ mutants have very low V-ATPase activity and assembly in vacuoles (Smardon et al., 2002). In this context, it was surprising that the SPVD chimera, which appears to lack Rav1 binding, had such high activity and assembly. We investigated the degree of RAVE independence of SPVD chimera by making a genomic deletion of *RAV1* in this mutant. The *rav1*Δ mutant grows well at pH 5 but poorly in media containing 4 mM Zn^2+^ or at elevated pH and calcium even at 30°C (Seol et al., 2001; Smardon et al., 2002). We found that the *rav1*ΔSPVD chimera grew as well as wild type at high pH and calcium concentration and quite well in zinc-containing medium at 30°C **(Figure 6)**. Thus, the growth defect of *rav1*ΔSPVD mutants is milder than *rav1*Δ mutants indicating that V-ATPases containing the SPVD chimera are at least somewhat independent of RAVE.

**Figure 6:**
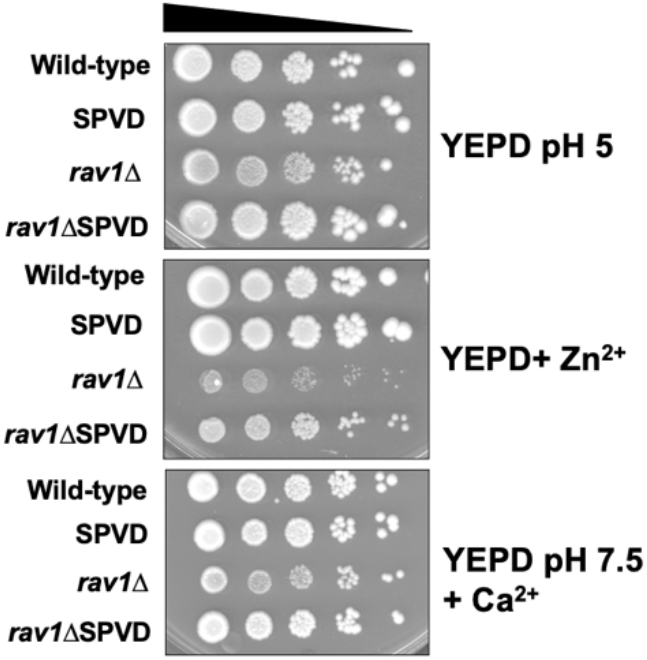
The SPVD chimera is partially independent of RAVE. Growth phenotypes of wild-type, SPVD, *rav1*Δ and *rav1*ΔSPVD strains were tested by growing strains to log phase in YEPD pH 5, diluting to an A_600_ of 0.8 and serially diluting 10-fold in a microtiter plate. Cells were transferred by pinning to YEPD pH 5 and YEPD + 4 mM ZnCl_2_ (YEPD +Zn^2+^) plates and grown for 3 days at 30°C. Cells transferred to YEPD pH 7.5 + 60 mM CaCl_2_ (YEPD pH 7.5 +Ca^2+^) plates were grown for 5 days at 30°C.

### Is there a physiological cost of the SPVD mutation?

Although V-ATPases containing the SPVD chimera appear to be fully assembled and functional, our *in vitro* experiments suggest that their regulatory properties may be quite different than those of wild-type V-ATPases. Reversible disassembly is a major regulatory mechanism for V-ATPases (Oot et al., 2017). In yeast, it is likely to be particularly important under conditions of acute glucose deprivation (Parra & Kane, 1998). We first checked whether V-ATPase regulation by reversible disassembly is retained in the SPVD chimera. We performed immunoprecipitations from whole cell lysates of wild-type and the SPVD chimera maintained in the presence of glucose, after 20 min of glucose deprivation, and 20 min after glucose readdition to the glucose-deprived cells. A monoclonal antibody against the V_1_ subunit B was previously shown to immunoprecipitate both V_1_ subcomplexes and fully assembled V_1_-V_o_ complexes (Doherty & Kane, 1993; Smardon et al., 2002). We used this antibody to immunoprecipitate complexes from both strains and assessed the ratio of V_1_ to V_o_ by comparing levels of V_1_ B-subunits to their respective V_o_ a-subunit, then calculating the percent assembly relative to the starting (+glucose) samples.

There are significant differences in the behavior of the two strains **(Figure 7A, B)**. V-ATPases in the SPVD mutant disassemble more extensively than those in the wild-type strain (a mean of 22% relative assembly vs. 51% in wild-type). Rapid reassociation of disassembled V_1_ and V_o_ after glucose restoration to glucose-deprived cells requires the RAVE complex *in vivo* (Seol et al., 2001). In wild type cells, the ratio of V_1_ to V_o_ was rapidly restored to a mean of 86% of the pre-deprivation level after glucose readdition **(Figure 7B)**. In the SPVD chimera the average assembly after glucose addition was less than 50% of the pre-deprivation level **(Figure 7B)**. Together, these data suggest that V-ATPases in SPVD chimera disassemble more extensively during glucose deprivation and cannot reassemble completely after a short time of glucose readdition.

**Figure 7:**
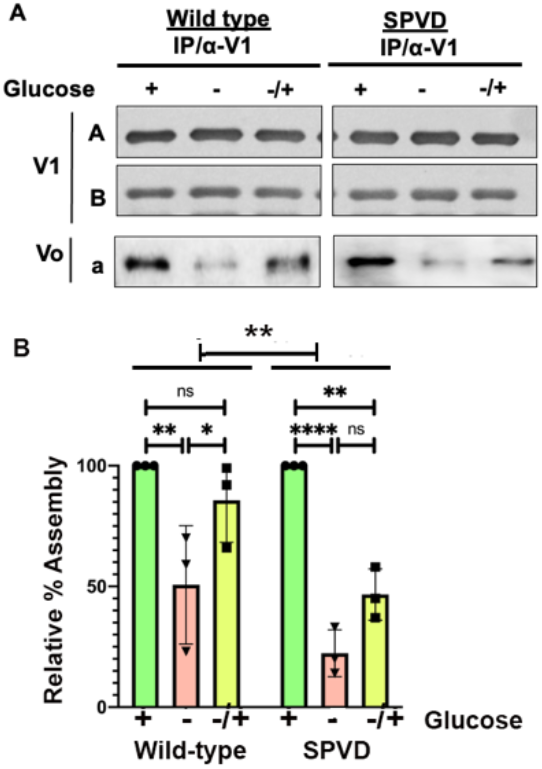
Glucose-dependent disassembly and reassembly in wild-type and SPVD cells. **A**. Immunoprecipitation of V-ATPases from wildtype and SPVD from spheroplast lysates was performed as described in (Kane, 1995) and Materials and Methods. Yeast cells were converted to spheroplasts, which were then incubated for 30 min in YEP media with (+) or without (-) 2% glucose. A third sample (-/+) was incubated without glucose for 20 min followed by 30 min incubation with 2% glucose. V-ATPases were immunoprecipitated under each condition using the monoclonal antibody 13D11 against the V_1_ B-subunit followed by Protein A-Sepharose. Immunoprecipitated proteins were subjected to SDS-PAGE and transferred to nitrocellulose. Western blot analysis was performed using primary antibody 8B1F3 (against the V_1_ A-subunit), 13D11 (against the V_1_ B-subunit), 7B1H1 (against Vph1NT) or polyclonal anti-Stv1 antisera. Dissociation of the V-ATPase complex is reflected as a decrease in the amount of a-subunit immunoprecipitated. A representative experiment from three biological replicates is shown. **B**. Bar diagram showing mean assembly under each condition, relative to the assembly each strain maintained in glucose of wild type and SPVD cells. Error bars represent standard deviation. Comparison between the strains was performed by two-way ANOVA (without multiple comparison). Results for +glucose, -glucose and glucose addback (-/+) conditions were also compared for each strain by two-way ANOVA with Tukey’s multiple comparison test. Significance of differences is indicated as: ****, P<0.0001; ***, P<0.0005; **, P<0.005; *, P<0.05; ns, P>0.05

Reversible disassembly of V-ATPases inhibits ATP hydrolysis and may support cell growth by preserving ATP during the periods of deprivation that arise during shifts in carbon source. We tested how cells containing the SPVD chimera adjust during a transition to a less preferred carbon source. Wild type and SPVD mutant cells were grown to log phase in YEPD (glucose) medium and diluted into fresh medium containing glucose or raffinose, then grown until growth reached a plateau. Both strains exhibited similar growth rates in medium containing 2% glucose as shown in **Figure 8A**, but the SPVD strain had a somewhat longer lag before growth initiation. There was also a lag in growth of wild-type cells upon a shift to raffinose; this is not unexpected because both transcriptome and proteome changes occur during a shift to raffinose metabolism (Paulo et al., 2015). SPVD cells have a significantly longer lag when shifted to raffinose than when shifted to fresh glucose-containing medium. These data indicate that with SPVD mutation cells cannot as readily adjust to changes in carbon sources as wild-type cells, consistent with the delayed V-ATPase reassembly in **Figure 8B**.

**Figure 8:**
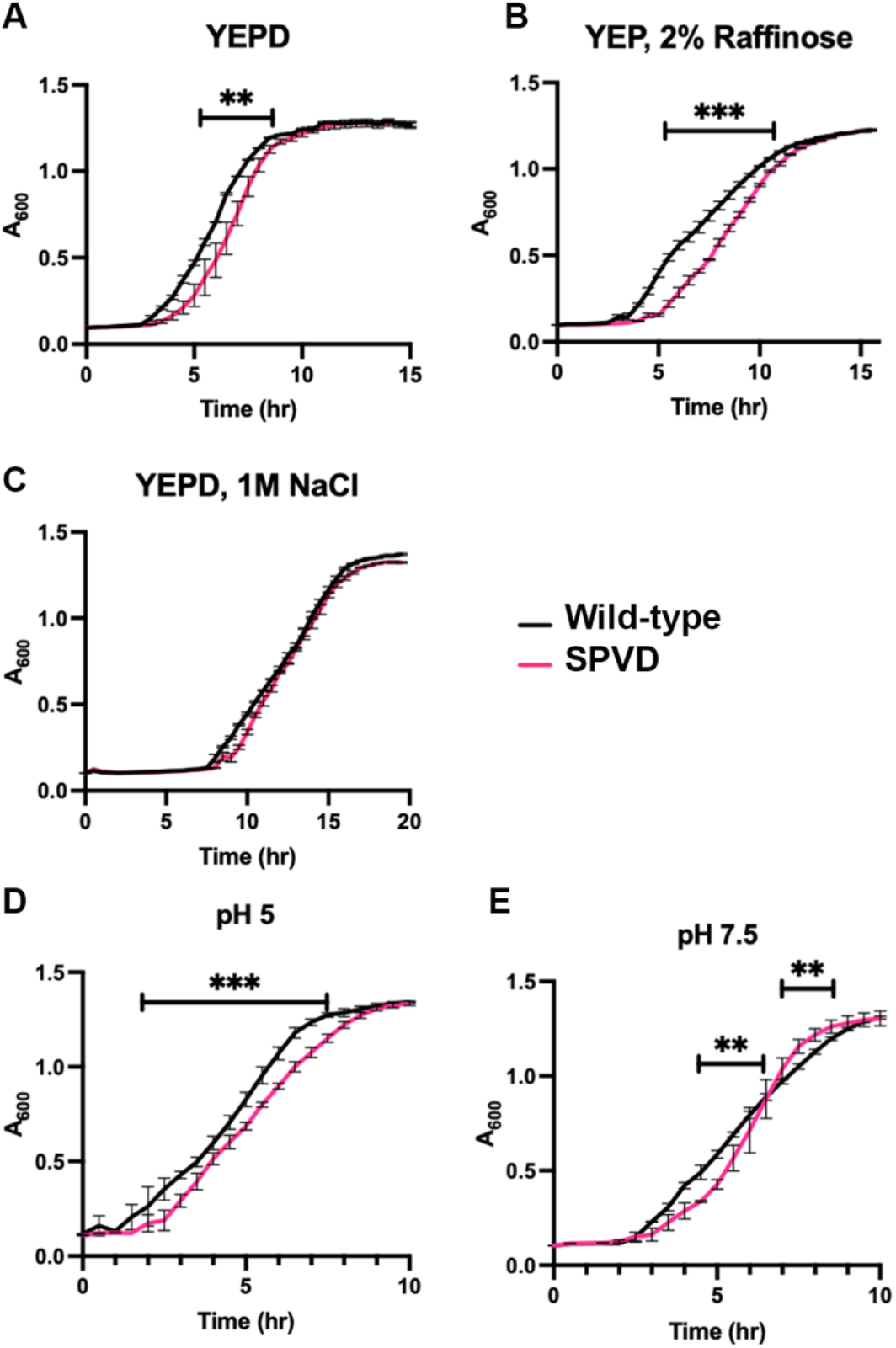
Growth of SPVD mutant under stress conditions. **A-C**. The wild-type and SPVD mutant strains were grown to log phase in YEPD, then diluted into the indicated medium to a density of A_600_=0.1. Growth was then monitored for 24 hr. at 30°C. Growth curves for wild type and SPVD strains in **A**. YEPD (YEP containing 2% glucose) **B**. 2% raffinose (YEP containing 2% raffinose) **C**. 1.0 M NaCl (YEPD containing 1 M NaCl) **D, E**. Wild-type and SPVD strains were grown to log phase in YEPD buffered to pH 5, then diluted into: **D**. pH 5 (YEPD buffered to pH 5) or **E**. pH 7.5 (YEPD buffered to pH 7.5). Growth was observed over the span of 24 hr at 30°C. Mean growth rates ± S.D. from three biological replicates are shown. Multiple t-tests were performed for each set of experiments. The bars show time ranges with significant differences in growth: ***, P<0.0005, **, P<0.005; all other differences were not significant.

PI(3,5)P_2_ is a signaling lipid that helps cells to adapt to multiple stresses (Jin et al., 2017; McCartney et al., 2014). Mutants lacking PI(3,5)P_2_ are sensitive to high salt (Bonangelino et al., 2002) and *vph1* mutants defective in PI(3,5)P_2_ activation also exhibit salt-dependent phenotypes (Banerjee et al., 2019; Bonangelino et al., 2002). We hypothesized that the SPVD chimera might have an advantage after a shift to medium containing high salt because it shows tighter binding to PI(3,5)P_2_. However, cells containing wild-type *VPH1* or the SPVD chimera showed very similar lags and growth rates **(Figure 8C)**. Under alkaline growth conditions, V-ATPase activity is increased and the enzyme appears to be stabilized (Diakov & Kane, 2010); this stabilization at alkaline pH is dependent on PI(3,5)P_2_ (Li et al., 2014). We hypothesized that increased PI(3,5)P_2_ binding of the SPVD chimera might provide a growth benefit when cells are shifted to alkaline pH (i.e. from medium at pH 5 to pH 7.5). The SPVD mutant again had a somewhat longer lag when diluted into pH 5 medium **(Figure 8D)**. However, at pH 7.5 the SPVD cells grew at a significantly faster rate than wild type cells after the initial lag (**Figure 8E**). These data indicate that the different regulatory interactions of the wild-type and SPVD aNT domains observed *in vitro* affect the response of cells to distinct stresses *in vivo*.

## DISCUSSION

The aNT domain occupies a critical location at the interface of the V_1_ and V_o_ subcomplexes and has long been recognized as a major locus of regulation. It is clear that different a-subunit isoforms endow V-ATPases with distinct regulatory capacities. However, <u>where</u> isoform-specific regulatory responses reside in the aNT domain and <u>how</u> cells decode these distinct regulatory signals are not completely understood.

The aNT chimeras presented here help define the hierarchy and consequences of isoform-specific regulation, as depicted in **Figure 9**. Specific binding of Vph1 to RAVE underlies the isoform-specific reversible disassembly of Vph1-containing V-ATPases (Smardon et al., 2014). Binding of the Rav1(678-898) fragment to the VPSD chimera in **Figure 2** indicates that the RAVE complex binds to the proximal end of Vph1NT (indicated by the white star in Figure 9.) The Rav1(678-898) fragment is critical for recruitment of RAVE to the vacuolar membrane; mutations in this region completely prevent recruitment (Jaskolka & Kane, 2020). Thus, the proximal end of Vph1NT may make the first contact between the V_o_ subcomplex and the RAVE complex bearing V_1_ subcomplex and subunit C. This initial contact will help orient the complexes for reassembly into the active complex (Jaskolka et al., 2021). Binding of RAVE to the proximal end of Vph1NT as shown for the wild-type V-ATPase in **Figure 9** could leave the distal end free to assemble a quaternary complex with the foot domain of subunit C and the appropriate EG peripheral stalk; formation of this quaternary interaction is likely to be a critical step in reassembly (Oot et al., 2017).

**Figure 9:**
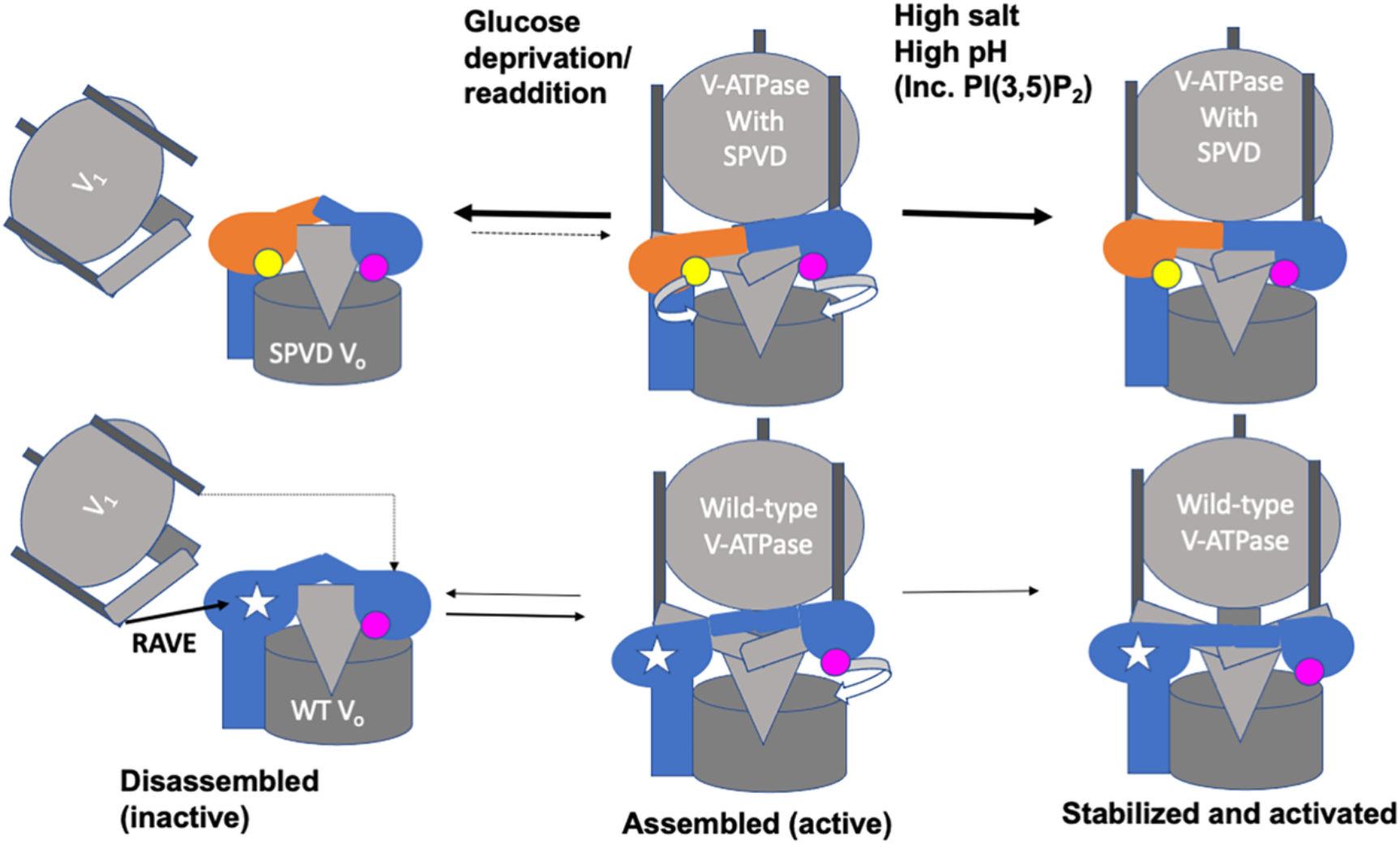
Model of regulatory properties of SPVD vs. wild-type V-ATPases. Both wild-type V-ATPases and V-ATPases containing the SPVD chimera are depicted as existing in three states depending on extracellular conditions. Based on data in Figure 7, the SPVD chimera disassembles more completely upon glucose deprivation and reassembles more slowly than wild-type upon glucose readdition. In contrast, based in Figure 8E, the SPVD chimera may reach a PI(3,5)P_2_-stabilized state more readily than the wild-type enzyme, allowing more rapid growth upon a shift to high pH. PIP binding sites are depicted as in Figure 1, and binding of the RAVE complex to the Vph1 proximal domain (which may assist during reassembly) is indicated by a white star.

The chimeric aNTs provide new insights into the mechanism of interactions of aNT isoforms with PIP lipids. Interestingly, lipid binding sites have been identified in poorly conserved loops of both the proximal domain (Stv1NT) (Banerjee & Kane, 2017) and the distal domain (Vph1NT) (Banerjee et al., 2019), suggesting both ends of aNT can potentially contribute to PIP binding and increased V-ATPase activity. We find that Vph1NT acquires PI(4)P binding *in vitro* upon addition of the previously defined WKY sequence from Stv1NT. However, both the SPVD chimera, which contains this sequence, and the VPSD chimera, which does not, show better binding to PI(4)P than Vph1NT, suggesting that other Stv1NT sequences contribute to PI(4)P binding. Previous attempts to determine direct interaction between Vph1NT and PI(3,5)P_2_ in a liposome flotation assay were unsuccessful, and weak binding was also observed here in the liposome pelleting assay **(Figure 3B,D)**. PI(3,5)P_2_ binding to Vph1NT is a reversible, regulatory interaction *in vivo* (Li et al., 2014), and thus may be tuned to a lower affinity. Interestingly, the SPVD chimera, which contains both of previously determined PIP binding sites, binds more tightly to <u>both</u> PI(4)P and PI(3,5)P_2_ than either Stv1NT or Vph1NT *in vitro*. Cryo-EM structures indicate that both the proximal and distal domains of the aNTs collapse toward the center of the disassembled, inactive V_o_ sector (Mazhab-Jafari et al., 2016; Roh et al., 2018). Lipid binding to either end of aNT could help stabilize the subunit in a more peripheral configuration optimal for assembly and activity. PIP binding to both ends of the SPVD chimera may provide even greater stability.

The SPVD chimera is the first fully functional chimera of Stv1 and Vph1. Kawasaki-Nishi et al. constructed chimeras containing the entire NT domain of Stv1 and Vph1 in combination with the CT domains of the opposite isoform (Kawasaki-Nishi et al., 2001). However, both of those chimeras had <30% of the activity and assembly than the wild-type enzyme, comparable to the 23% of wild-type activity in the VPSD mutant (**Figure 5A**). In contrast, the assembly and activity of the SPVD chimera in isolated vacuolar vesicles are at least as good as those of the wild-type enzyme. This indicates that binding of the RAVE complex to the proximal end of the chimeric aNT is either not essential for the biosynthetic assembly V-ATPases containing the SPVD chimera, or that the proximal end of Stv1NT is sufficient to support RAVE-independent biosynthesis. Notably, Stv1-containing V-ATPases can assemble in the absence of RAVE (Smardon et al., 2014), so this capability may be carried in the Stv1NT proximal end. However, it is also possible that the increased lipid binding to SPVD-containing ATPases circumvents the need for RAVE in biosynthetic assembly. In addition to its biosynthetic role, RAVE is required for reassembly after glucose deprivation (Seol et al., 2001; Smardon et al., 2002). The SPVD chimera disassembles to a greater extent than the wild-type V-ATPase and incompletely reassembles in a short time after glucose readdition, as shown in **Figure 8** and depicted in **Figure 9**. Biosynthetic assembly and reassembly occur by different pathways, so RAVE interactions may differ between the two pathways (Kane, 2006).

The SPVD chimera provides novel insights into how isoform-specific regulatory inputs are integrated and balanced to achieve physiological responses (**Figure 9**). PIP binding and reversible disassembly are associated with V-ATPase responses to different stresses (Kane, 1995; Li et al., 2012), and their regulatory effects are separable. Reversible disassembly in response to glucose deprivation still occurs in the absence of vacuolar PI(3,5)P_2_, suggesting that this process is independent of PIP binding (Li et al., 2014). However, the V-ATPase response to glucose restoration is strongly RAVE-dependent (Seol et al., 2001). V-ATPase disassembly can occur with acute glucose deprivation, from exhaustion of glucose during growth, and in transitions to poor carbon sources (Kane, 1995; Parra & Kane, 1998). We found that despite having wild-type activity in isolated vacuoles, the SPVD mutant was slow to resume growth after a shift from glucose to the poor carbon source raffinose **(Figure 8B**). In other physiological situations, RAVE binding may be less important. Specifically, vacuolar PI(3,5)P_2_ is implicated in stabilization and/or activation of assembled V-ATPases at the vacuole under high salt and alkaline growth conditions (Diakov & Kane, 2010; Li et al., 2014). PI(3,5)P_2_ stabilization of V-ATPases under alkaline stress could help explain the higher rate of recovery of the SPVD mutant at pH 7.5, after an initial lag (**Figure 8E**). Further experiments will be needed to fully define the underlying mechanism, but this is the first example of a chimeric V-ATPase with both preserved basal activity and a growth advantage over wild-type under specific stress conditions.

Total loss of V-ATPase activity is lethal in higher eukaryotes (Sun-Wada et al., 2003), so targeting isoform-specific functions and regulatory mechanisms is an attractive route to therapeutic modulation of V-ATPase activity. Generation of functional V-ATPases with altered regulatory properties provides a means to better understand both the mechanism and underlying balance of isoform-specific regulation. Importantly, our results indicate that it may be possible to retain full V-ATPase activity under basal conditions, thus avoiding lethality, while changing the enzyme’s regulation and cellular responses to stress.

## MATERIALS AND METHODS

### Materials, media and cell growth

Amylose and TALON resin were purchased from New England Biolabs and Clontech, respectively. Anti-FLAG M2 resin, mouse anti-FLAG antibody, and FLAG peptide were purchased from Sigma. Lipids were purchased as lyophilized powder from Avanti Polar Lipids. Sepharose A beads were purchased from GE Healthcare. DSP (dithiobis (succinimidyl propionate)) crosslinker was obtained from Pierce.

Liquid medium for E. coli culture consisted of 2.5% Luria–Bertani broth (LB), Miller powder (Fisher BioReagents). For expression of proteins from E. coli 0.2% dextrose was added to LB. To maintain plasmids carrying an ampicillin- or chloramphenicol resistance genes, we added 125 μg/ml ampicillin (Sigma-Aldrich) or 34 μg/ml chloramphenicol (Sigma-Aldrich), respectively.

Yeast cells were grown on rich media containing 1% yeast extract, 2% peptone and 2% dextrose (YEPD). YEPD was buffered to pH 5.0 or pH 7.5 by 50 mM potassium phosphate and 50 mM potassium succinate. For plates with added CaCl_2_, YEPD was buffered to pH 7.5 with 50 mM Mes (2-(N-morpholino) ethanesulfonic acid) 50 mM MOPS (3-(N-morpholino) propanesulfonic acid) buffer, and 60 mM CaCl_2_ was added (YEPD, pH 7.5, Ca^2+^). YEPD+Zn^2+^ containing plates consisted of 4 mM ZnCl_2_ in unbuffered YEPD. YEPD+G418 contained 200 μg/ml G418 (Gibco Genticin, Thermo Fisher Scientific). For auxotrophic selections, fully supplemented minimal medium with 2% dextrose (SD) lacking individual nutrients was used. Yeast cells were grown to log phase in indicated media and collected near A_600_=1 for transformation as described (Gietz & Schiestl, 2007). For dilution growth assays, the indicated wild-type or mutant yeast cells were grown to log phase in YEPD, pH 5. Cell suspensions were adjusted to a single density, and 10-fold serial dilutions were made and then pinned onto the desired plates.

### Yeast strains

The genotypes of the strains used in this study are listed in Table 1.

**Table 1:**
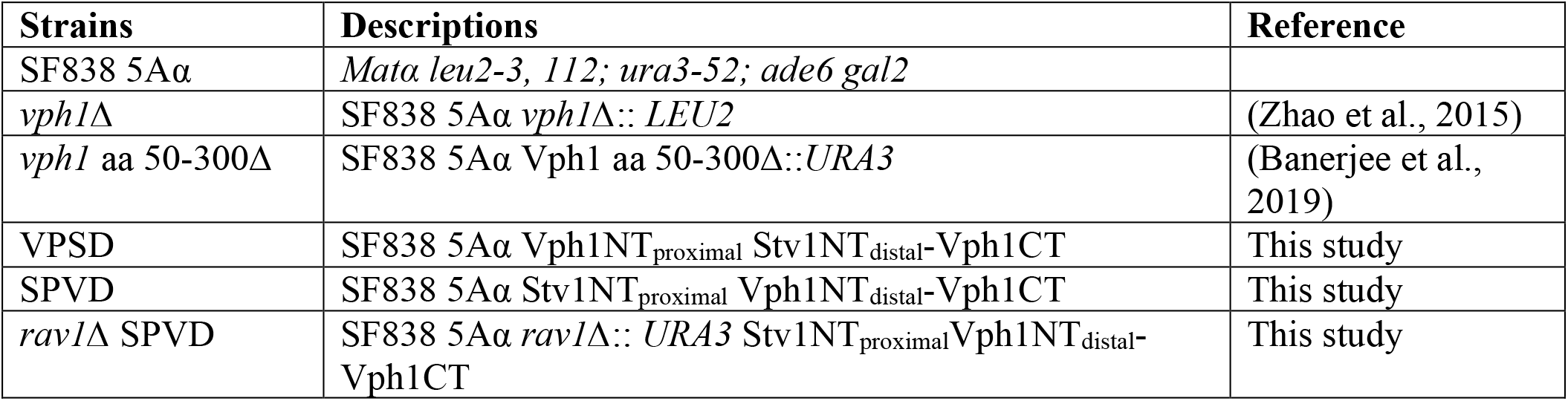
Yeast strains used in this study.

For genomic integration of the VPSD chimeric aNT into full length *VPH1* under the *VPH1* promoter, a fragment containing VPSD chimera was PCR amplified from the expression plasmid described above using oligonucleotides SIV F and SIV R. This amplified PCR product was then transformed into a strain in which Vph1NT sequence for amino acids 50-300 had been replaced with *URA3* (Banerjee et al., 2019). Transformants capable of growth in presence of 5-fluoro-orotic acid (FOA) were selected, analyzed for replacement of the *URA3* gene, and sequenced to confirm introduction of the chimera sequence.

For genomic integration of SPVD chimeric aNT into full length *VPH1* under the *VPH1* promoter, a plasmid containing 400 bp upstream of the *VPH1* open reading frame followed by the SPVD chimera and 400 bp corresponding to Vph1CT amino acids 406-539 cloned into pBluescript II KS(-) vector using BamHI and XhoI was purchased from Genscript. After restriction digestion with BamH1 and Xho1, the fragment was then transformed into 5Aα Vph1 aa 50-300Δ::*URA3* strain and transformants were selected on FOA plates as described above.

To delete *RAV1* from the SPVD-containing yeast strain, nucleotide sequence of *URA3* with ends homologous to *RAV1* was PCR amplified from a strain containing *rav1*Δ::*URA3* strain using oligonucleotides rav1 Ura3F and Rav1 Ura3R. This amplified fragment was used to transform 5Aα SPVD cells and transformants were selected on SD lacking uracil. Colony PCR was performed to isolate transformants with the *RAV1* deletion.

### Cloning, expression and purification of MBP-Vph1NT(1-372)-FLAG, VPSD chimera, SPVD chimera and Vph1NT-WKY mutant

MBP-Vph1NT(1-372)-FLAG: A single FLAG tag was inserted at the C-terminus of MBP-Vph1NT (1-372) by TA cloning. Specifically, Vph1NT nucleotide sequence 1-1116 carrying a 5’ BamH1 and a 3’ FLAG was PCR amplified using oligonucleotides Vph1NT 5’BamH1 and Vph1NT 3’FLAG (Table 2) and a template of MBP-Vph1NT (1-372) plasmid DNA. After addition of a non-templated A, the PCR product was then ligated in pGEM T-easy vector. Subsequently, BamH1-Vph1NT-FLAG was cloned into pMal c2E Precision protease vector that had been cut with restriction enzymes BamHI and PstI.

**Table 2:**
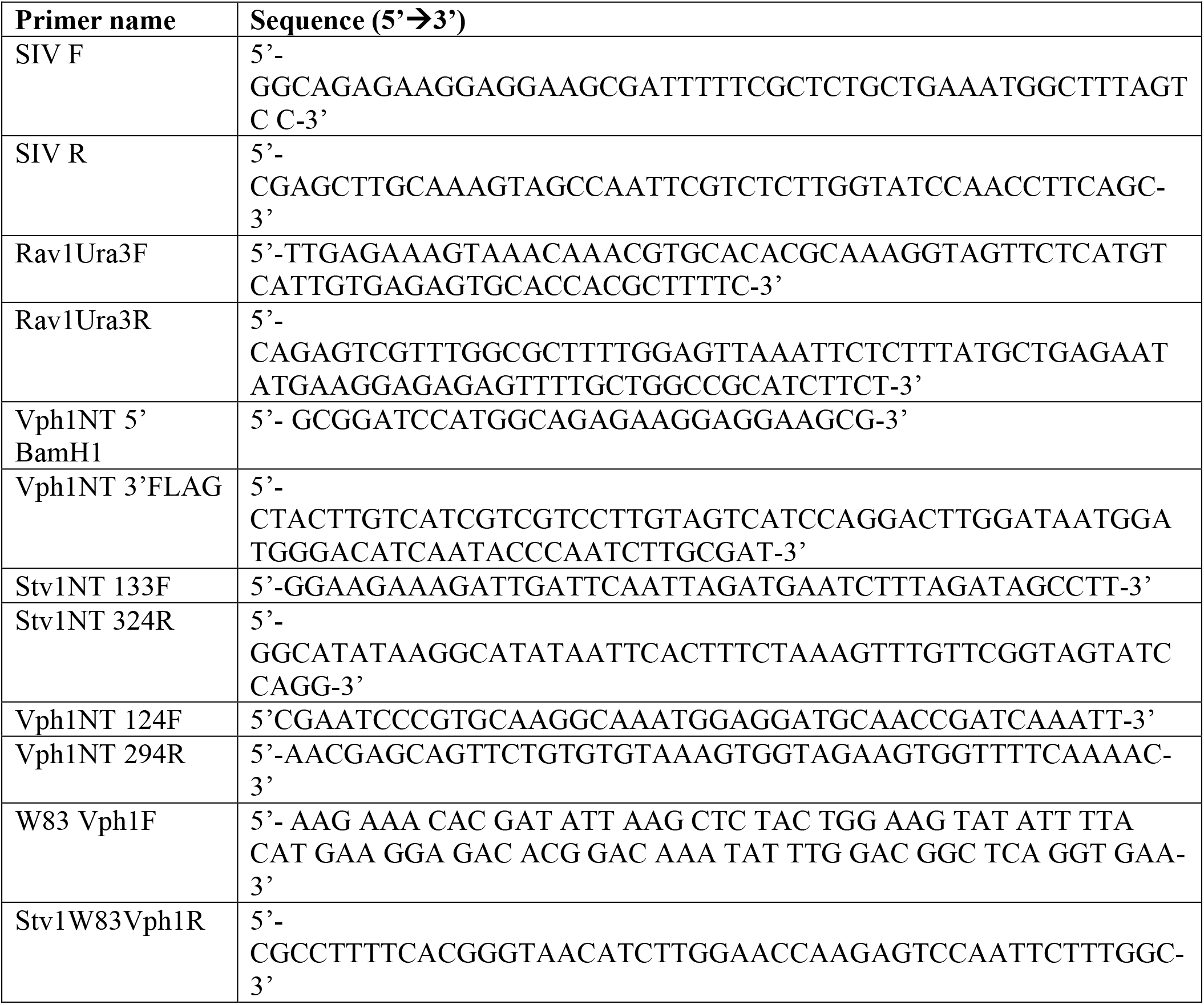
Oligonucleotides used in this study.

The VPSD chimera carrying amino acid residues 1-123 of Vph1NT, 133-342 of Stv1NT and 295-372 of Vph1NT was generated by using a megaprimer PCR technique (Forloni et al., 2019). A sequence of Vph1NT corresponding to amino acid 124-294 was replaced with sequence corresponding to Stv1NT amino acid 133-342 using oligonucleotides Stv1NT 133F and Stv1NT 324R and a template of MBP-Vph1NT (1-372)-FLAG plasmid DNA.

The SPVD chimera carrying amino acid residues 1-132 of Stv1NT, 124-294 of Vph1NT and 343-418 of Stv1NT was generated by using similar megaprimer PCR technique. A sequence of Stv1NT corresponding to amino acid 133-342 was replaced with sequence corresponding to Vph1NT amino acid 124-294 using oligonucleotides Vph1NT 124F and Vph1NT 294R and a template of MBP-Stv1NT(1-418)-FLAG plasmid DNA.

For the Vph1NT-WKY mutant, a mutant Vph1NT carrying Stv1NT W83KYILH amino acid was generated by using oligonucleotide W83 Vph1F and Stv1W83Vph1R and a template of MBP-Vph1NT (1-372)-FLAG plasmid DNA.

MBP-Vph1NT(1-372)-FLAG, MBP-Stv1NT(1-418)-FLAG, VPSD chimera, SPVD chimera, Vph1NT-WKY mutant and MBP-Rav1 (679-898)-His_6_ cell were expressed, induced and affinity purified on amylose columns as described previously (Jaskolka & Kane, 2020). All the constructs were expressed in Rosetta competent cells (F^-^ *ompT hsdS*_B_(r_B_^-^m_B_^-^) *gal dcm* (DE3) pRARE (Cam^R^)) (Novagen). To induce protein expression 0.5mM isopropyl B-D-thiogalactopyranoside was added to cells when they reached a density of 0.5-0.6 OD/ml, and growth was continued for 16 hr at 18°C. Proteins were affinity purified on amylose columns. A second round of affinity purification was performed using anti-FLAG M2 Affinity resin. Peak fractions were pooled, and 5 mM DTT was added immediately. After concentration to at least 1 mg/ml, the proteins were run on a Sephadex 200 (S200) gel-filtration fast protein liquid chromatography (FPLC) column on an AKTA FPLC system. The peak corresponding to the monomeric molecular mass was collected and used for additional experiments.

### Pull-down assay

Protein pull-down assays were performed as described (Jaskolka & Kane, 2020). The only exception is that the two step affinity purified peak fractions of MBP-Vph1NT(1-372)-FLAG, MBP-Stv1NT(1-418)-FLAG, VPSD chimera, SPVD chimera and pooled peak fractions of Rav1 (679–898)-His_6_ from amylose column were dialyzed overnight in TALON equilibration buffer. MBP was cleaved from the proteins with Prescission protease before affinity purification on the FLAG column. All four aNT proteins were brought to a similar concentration (∼1 μM). The aNT proteins were each mixed with the Rav1 (679-898)His_6_ at a 5:1 molar ratio. The mixtures were added to TALON resin and incubated at 4°C for 2 h. to allow binding, then washed and separated by SDS-PAGE.

### Liposome preparation and liposome pelleting assay

Control liposomes and liposomes of defined PIP lipid contents are prepared as described (Banerjee & Kane, 2017). For pelleting assays, the protein of interest was mixed with a final volume of 100 μl of control (0% PIP) or experimental (5% PIP) liposomes to give a final concentration of protein of 0.5 or 1 μM, and final total lipid concentration of 0.33 mM. The mixtures were left at room temperature for 30 min to allow for protein–liposome interaction. After incubation, the solution was centrifuged at 400,000 g for 30 min at 4°C in a TLA-100 fixed angle rotor using a Beckman Coulter mini-ultracentrifuge. Supernatant and pellet fractions were separated, and the pellet was resuspended in 100 μl of 50 mM Tris-Cl, 100 mM NaCl (pH 7.4). Samples were then precipitated using 10% trichloroacetic acid. The pellets were washed with cold acetone and dissolved in 50 μl of cracking buffer (50 mM Tris-HCl, pH 6.8, 8 M urea, 5% SDS, and 1 mM EDTA). 25 μl of each sample were then separated by SDS-PAGE and transferred into nitrocellulose membrane. The resulting blots were then probed with anti-FLAG M2 antibody, followed by alkaline phosphatase-conjugated anti-mouse antibody. A no liposome control was also performed to ensure that the protein of interest did not pellet on its own. For these samples, the protein was diluted to final volume of 100 μl in the same buffer and subjected to centrifugation and SDS-PAGE as described above.

A least three separate pelleting assays were performed for each protein concentration and liposome type. Immunoblots were analyzed using ImageJ to quantitate intensity of the supernatant and pellet bands which was subsequently converted to percentage of the total (supernatant + pellet). For each experiment, the percentage of protein in the control (no PIP) pellets were subtracted from that in the PIP pellets and the resulting value of specific binding was multiplied by the total protein concentration to determine the PIP-specific binding. The mean and standard deviation across replicates were calculated, and significance was determined using Ordinary one-way ANOVA with multiple comparisons using GraphPad Prism 9.

### Vacuole isolation and biochemical analysis

Vacuoles were isolated as described in (Li et al., 2014). Purified vacuolar vesicles were tested for ATPase activity in a coupled ATPase enzyme assay as described in (Liu et al., 2005) except that the cuvettes containing reaction mixture were preheated at 37°C. To assess V-ATPase specific activity, concanamycin A was added to a final concentration of 200 nM. Protein concentration was determined by Lowry assay (Lowry et al., 1951) to determine V-ATPase specific activity. Proton pumping was observed using the 9-amino-6-chloro-2-methoxyacridine quenching assay described previously (Liu et al., 2005). 20 μg of vacuolar vesicles were used for each assay. Pumping was initiated by adding 0.5 mM ATP and 1.0 mM MgSO4.

Western blots of vacuolar proteins were performed as in (Li et al., 2014). Mouse monoclonal antibodies 8B1F3 and13D11 were used against V_1_ subunits A and B (Kane et al., 1992). Vph1 was detected using mouse monoclonals 7B1H1 and 10D7 (Kane et al., 1992). Alkaline phosphatase was detected with mouse monoclonal 1D3A10 (Invitrogen). Rabbit polyclonal antisera, a generous gift from Tom Stevens, University of Oregon, were used to detect V_1_ C and V_o_ d and SPVD (via anti-Stv1p antisera). Representative immunoblots were scanned to obtain images used for figures.

### Immunoprecipitation

Immunoprecipitation was performed as described in (Kane, 1995) with the following modifications. For each immunoprecipitation, volumes of cell culture equivalent to 20 OD_600_ units were pelleted then converted to spheroplasts. The spheroplasts were incubated for 30 min in YEP, 1.2 M sorbitol media with (+) or without (-) 2% glucose at 30°C. A third sample (-/+) was incubated without glucose for 20 min followed by 30 min incubation with 2% glucose. The spheroplasts were lysed, then complexes immunoprecipitated using the monoclonal antibody 13D11 against the B-subunit followed by Protein A Sepharose beads (Sigma). After elution with cracking buffer, (8M urea, 50mM Tris HCL pH 6.8, 5% SDS, 1mM EDTA, 5% β-mercaptoethanol) samples were vortexed and incubated for 20 min at 55°C. All samples were then subjected to SDS-PAGE and transferred to nitrocellulose. Western blot analysis was performed using primary antibody 8B1F3 (V_1_ A-subunit), 13D11 (V_1_ B-subunit), 7B1H1 (Vph1 in wild type) and anti-Stv1 antibody (SPVD). Alkaline phosphatase conjugate secondary antibody was used for the detection of Vph1 in wild type whereas HRP conjugate secondary antibody was used for the detection of the SPVD a-subunit. The extent of V-ATPase assembly was assessed from the ratio of V_o_ to V_1_ and normalized to ratio in the sample maintained in glucose.

### Growth curve assay

For growth assays, yeast strains were grown overnight in YEPD and diluted to a density of A_600_ = 0.05 in YEP containing 2% dextrose or 2% raffinose or YEPD containing 1 M NaCl. In order to compare pH effects on growth, cells were grown overnight in YEPD buffered to pH 5, then diluted to YEPD buffered to pH 5 or pH 7.5 as indicated. Growth over time was then monitored on a SpectraMax i3X multi-mode multiplate reader for 24 hr.

### Protein structure prediction

The homology models of Vph1NT (1-372) and Stv1NT (1-448) were obtained by submitting the corresponding amino acid sequences of Phyre2 server (Kelley et al., 2015). From the submitted sequence, 100% confidence models based on Vph1NT in the state 1 rotational structure of the yeast V-ATPase (Zhao et al., 2015) (PDB 3j9t) were obtained with 93% coverage (Vph1NT 1-372) and 88% coverage (Stv1NT 1-418). Structures were visualized using UCSF Chimera (Pettersen et al., 2004).

### Statistical analysis

Statistical analysis was performed using Graphpad Prism 9. For every experiment a minimum of three independent experiments was performed. In the graphs in Figures 3, 5, and 7, each dot superimposed on the bar graphs represents a biological replicate. Biological replicates are defined as coming from distinct individual yeast colonies on a plate (in contrast, experiments in which a liquid culture of yeast is grown and divided would give rise to technical replicates). For experiments with purified protein, biological replicates come from different purifications of a given protein. All datasets were analyzed for outliers using Prism 9; only one outlier (which had very high activity) was identified and excluded in the vacuolar ATPase assays from the SPVD mutant (Figure 5A).

## Abbreviations

V-ATPase: V-type proton-translocating ATPase
aNT: N-terminal cytosolic domain of the V-ATPase a-subunit
SPVD: chimeric aNT containing the proximal domain of Stv1 and distal domain of Vph1
VPSD: chimeric aNT containing the proximal domain of Vph1 and distal domain of Stv1
RAVE: regulator of acidification of vacuoles and endosomes.

## ACKNOWLEDGEMENTS

This work was supported by NIH R01 GM126020 to P.M.K. The authors thank Juliana Bourgeios and Maureen Tarsio for excellent technical help, Dr. Tom Stevens (University of Oregon) for antibodies, and Drs. Stephan Wilkens and Rebecca Oot (Upstate Medical University) for helpful discussion and use of the FPLC.

## COMPETING INTERESTS

The authors declare no competing interests.

## Notes

### Competing Interest Statement

The authors have declared no competing interest.

### Summary of Updates

Revisions have been made to Figures 1,2,3,4 and Supplemental Figure 1 and a new Figure 9 model has been added. Corresponding changes to the text have also been made.

## REFERENCES

Banerjee, S., Clapp, K., Tarsio, M., & Kane, P. M. (2019). Interaction of the late endo-lysosomal lipid PI(3,5)P2 with the Vph1 isoform of yeast V-ATPase increases its activity and cellular stress tolerance. J Biol Chem, 294(23), 9161–9171. https://doi.org/10.1074/jbc.RA119.008552

Banerjee, S., & Kane, P. M. (2017). Direct interaction of the Golgi V-ATPase a-subunit isoform with PI(4)P drives localization of Golgi V-ATPases in yeast. Mol Biol Cell, 28(19), 2518–2530. https://doi.org/10.1091/mbc.E17-05-0316

Benlekbir, S., Bueler, S. A., & Rubinstein, J. L. (2012). Structure of the vacuolar-type ATPase from Saccharomyces cerevisiae at 11-A resolution. Nat Struct Mol Biol, 19(12), 1356–1362. https://doi.org/10.1038/nsmb.2422

Bonangelino, C. J., Nau, J. J., Duex, J. E., Brinkman, M., Wurmser, A. E., Gary, J. D., Emr, S. D., & Weisman, L. S. (2002). Osmotic stress-induced increase of phosphatidylinositol 3,5-bisphosphate requires Vac14p, an activator of the lipid kinase Fab1p. J Cell Biol, 156(6), 1015–1028. https://doi.org/10.1083/jcb.200201002

Breton, S., & Brown, D. (2013). Regulation of luminal acidification by the V-ATPase. Physiology (Bethesda), 28(5), 318–329. https://doi.org/10.1152/physiol.00007.2013

Casey, J. R., Grinstein, S., & Orlowski, J. (2010). Sensors and regulators of intracellular pH. Nat Rev Mol Cell Biol, 11(1), 50–61. https://doi.org/10.1038/nrm2820

Chandra, M., Chin, Y. K., Mas, C., Feathers, J. R., Paul, B., Datta, S., Chen, K. E., Jia, X., Yang, Z., Norwood, S. J., Mohanty, B., Bugarcic, A., Teasdale, R. D., Henne, W. M., Mobli, M., & Collins, B. M. (2019). Classification of the human phox homology (PX) domains based on their phosphoinositide binding specificities. Nat Commun, 10(1), 1528. https://doi.org/10.1038/s41467-019-09355-y

Collins, M. P., Stransky, L. A., & Forgac, M. (2020). AKT Ser/Thr kinase increases V-ATPase-dependent lysosomal acidification in response to amino acid starvation in mammalian cells. J Biol Chem, 295(28), 9433–9444. https://doi.org/10.1074/jbc.RA120.013223

Couoh-Cardel, S., Milgrom, E., & Wilkens, S. (2015). Affinity Purification and Structural Features of the Yeast Vacuolar ATPase Vo Membrane Sector. J Biol Chem, 290(46), 27959–27971. https://doi.org/10.1074/jbc.M115.662494

Diakov, T. T., & Kane, P. M. (2010). Regulation of vacuolar proton-translocating ATPase activity and assembly by extracellular pH. J Biol Chem, 285(31), 23771–23778. https://doi.org/10.1074/jbc.M110.110122

Doherty, R. D., & Kane, P. M. (1993). Partial assembly of the yeast vacuolar H(+)-ATPase in mutants lacking one subunit of the enzyme. J Biol Chem, 268(22), 16845–16851. https://www.ncbi.nlm.nih.gov/pubmed/8344963

Eaton, A. F., Merkulova, M., & Brown, D. (2021). The H(+)-ATPase (V-ATPase): from proton pump to signaling complex in health and disease. Am J Physiol Cell Physiol, 320(3), C392–C414. https://doi.org/10.1152/ajpcell.00442.2020

Finnigan, G. C., Cronan, G. E., Park, H. J., Srinivasan, S., Quiocho, F. A., & Stevens, T. H. (2012). Sorting of the yeast vacuolar-type, proton-translocating ATPase enzyme complex (V-ATPase): identification of a necessary and sufficient Golgi/endosomal retention signal in Stv1p. J Biol Chem, 287(23), 19487–19500. https://doi.org/10.1074/jbc.M112.343814

Forgac, M. (2007). Vacuolar ATPases: rotary proton pumps in physiology and pathophysiology. Nat Rev Mol Cell Biol, 8(11), 917–929. https://doi.org/10.1038/nrm2272

Forloni, M., Liu, A. Y., & Wajapeyee, N. (2019). Megaprimer Polymerase Chain Reaction (PCR)-Based Mutagenesis. Cold Spring Harb Protoc, 2019(6). https://doi.org/10.1101/pdb.prot097824

Frattini, A., Orchard, P. J., Sobacchi, C., Giliani, S., Abinun, M., Mattsson, J. P., Keeling, D. J., Andersson, A. K., Wallbrandt, P., Zecca, L., Notarangelo, L. D., Vezzoni, P., & Villa, A. (2000). Defects in TCIRG1 subunit of the vacuolar proton pump are responsible for a subset of human autosomal recessive osteopetrosis. Nat Genet, 25(3), 343–346. https://doi.org/10.1038/77131

Gietz, R. D., & Schiestl, R. H. (2007). High-efficiency yeast transformation using the LiAc/SS carrier DNA/PEG method. Nat Protoc, 2(1), 31–34. https://doi.org/10.1038/nprot.2007.13

Huotari, J., & Helenius, A. (2011). Endosome maturation. EMBO J, 30(17), 3481–3500. https://doi.org/10.1038/emboj.2011.286

Hurtado-Lorenzo, A., Skinner, M., El Annan, J., Futai, M., Sun-Wada, G. H., Bourgoin, S., Casanova, J., Wildeman, A., Bechoua, S., Ausiello, D. A., Brown, D., & Marshansky, V. (2006). V-ATPase interacts with ARNO and Arf6 in early endosomes and regulates the protein degradative pathway. Nat Cell Biol, 8(2), 124–136. https://doi.org/10.1038/ncb1348

Jaskolka, M. C., & Kane, P. M. (2020). Interaction between the yeast RAVE complex and Vph1-containing Vo sectors is a central glucose-sensitive interaction required for V-ATPase reassembly. J Biol Chem. https://doi.org/10.1074/jbc.RA119.011522

Jaskolka, M. C., Winkley, S. R., & Kane, P. M. (2021). RAVE and Rabconnectin-3 Complexes as Signal Dependent Regulators of Organelle Acidification. Front Cell Dev Biol, 9, 698190. https://doi.org/10.3389/fcell.2021.698190

Jin, N., Jin, Y., & Weisman, L. S. (2017). Early protection to stress mediated by CDK-dependent PI3,5P2 signaling from the vacuole/lysosome. J Cell Biol, 216(7), 2075–2090. https://doi.org/10.1083/jcb.201611144

Kane, P. M. (1995). Disassembly and reassembly of the yeast vacuolar H(+)-ATPase in vivo. J Biol Chem, 270(28), 17025–17032. https://www.ncbi.nlm.nih.gov/pubmed/7622524

Kane, P. M. (2006). The where, when, and how of organelle acidification by the yeast vacuolar H+-ATPase. Microbiol Mol Biol Rev, 70(1), 177–191. https://doi.org/10.1128/MMBR.70.1.177-191.2006

Kane, P. M. (2012). Targeting reversible disassembly as a mechanism of controlling V-ATPase activity. Curr Protein Pept Sci, 13(2), 117–123. https://www.ncbi.nlm.nih.gov/pubmed/22044153

Kane, P. M., Kuehn, M. C., Howald-Stevenson, I., & Stevens, T. H. (1992). Assembly and targeting of peripheral and integral membrane subunits of the yeast vacuolar H(+)-ATPase. J Biol Chem, 267(1), 447–454. https://www.ncbi.nlm.nih.gov/pubmed/1530931

Kawasaki-Nishi, S., Bowers, K., Nishi, T., Forgac, M., & Stevens, T. H. (2001). The amino-terminal domain of the vacuolar proton-translocating ATPase a subunit controls targeting and in vivo dissociation, and the carboxyl-terminal domain affects coupling of proton transport and ATP hydrolysis. J Biol Chem, 276(50), 47411–47420. https://doi.org/10.1074/jbc.M108310200

Kelley, L. A., Mezulis, S., Yates, C. M., Wass, M. N., & Sternberg, M. J. (2015). The Phyre2 web portal for protein modeling, prediction and analysis. Nat Protoc, 10(6), 845–858. https://doi.org/10.1038/nprot.2015.053

Li, S. C., Diakov, T. T., Rizzo, J. M., & Kane, P. M. (2012). Vacuolar H+-ATPase works in parallel with the HOG pathway to adapt Saccharomyces cerevisiae cells to osmotic stress. Eukaryot Cell, 11(3), 282–291. https://doi.org/10.1128/EC.05198-11

Li, S. C., Diakov, T. T., Xu, T., Tarsio, M., Zhu, W., Couoh-Cardel, S., Weisman, L. S., & Kane, P. M. (2014). The signaling lipid PI(3,5)P(2) stabilizes V(1)-V(o) sector interactions and activates the V-ATPase. Mol Biol Cell, 25(8), 1251–1262. https://doi.org/10.1091/mbc.E13-10-0563

Liu, M., Tarsio, M., Charsky, C. M., & Kane, P. M. (2005). Structural and functional separation of the N- and C-terminal domains of the yeast V-ATPase subunit H. J Biol Chem, 280(44), 36978–36985. https://doi.org/10.1074/jbc.M505296200

Lowry, O. H., Rosebrough, N. J., Farr, A. L., & Randall, R. J. (1951). Protein measurement with the Folin phenol reagent. J Biol Chem, 193(1), 265–275. https://www.ncbi.nlm.nih.gov/pubmed/14907713

Lu, M., Ammar, D., Ives, H., Albrecht, F., & Gluck, S. L. (2007). Physical interaction between aldolase and vacuolar H+-ATPase is essential for the assembly and activity of the proton pump. J Biol Chem, 282(34), 24495–24503. https://doi.org/10.1074/jbc.M702598200

Manolson, M. F., Proteau, D., Preston, R. A., Stenbit, A., Roberts, B. T., Hoyt, M. A., Preuss, D., Mulholland, J., Botstein, D., & Jones, E. W. (1992). The VPH1 gene encodes a 95-kDa integral membrane polypeptide required for in vivo assembly and activity of the yeast vacuolar H(+)-ATPase. J Biol Chem, 267(20), 14294–14303. https://www.ncbi.nlm.nih.gov/pubmed/1385813

Manolson, M. F., Wu, B., Proteau, D., Taillon, B. E., Roberts, B. T., Hoyt, M. A., & Jones, E. W. (1994). STV1 gene encodes functional homologue of 95-kDa yeast vacuolar H(+)-ATPase subunit Vph1p. J Biol Chem, 269(19), 14064–14074. https://www.ncbi.nlm.nih.gov/pubmed/7514599

Mazhab-Jafari, M. T., Rohou, A., Schmidt, C., Bueler, S. A., Benlekbir, S., Robinson, C. V., & Rubinstein, J. L. (2016). Atomic model for the membrane-embedded VO motor of a eukaryotic V-ATPase. Nature, 539(7627), 118–122. https://doi.org/10.1038/nature19828

McCartney, A. J., Zhang, Y., & Weisman, L. S. (2014). Phosphatidylinositol 3,5-bisphosphate: low abundance, high significance. Bioessays, 36(1), 52–64. https://doi.org/10.1002/bies.201300012

Oot, R. A., Couoh-Cardel, S., Sharma, S., Stam, N. J., & Wilkens, S. (2017). Breaking up and making up: The secret life of the vacuolar H(+) -ATPase. Protein Sci, 26(5), 896–909. https://doi.org/10.1002/pro.3147

Oot, R. A., & Wilkens, S. (2012). Subunit interactions at the V1-Vo interface in yeast vacuolar ATPase. J Biol Chem, 287(16), 13396–13406. https://doi.org/10.1074/jbc.M112.343962

Parra, K. J., & Kane, P. M. (1998). Reversible association between the V1 and V0 domains of yeast vacuolar H+-ATPase is an unconventional glucose-induced effect. Mol Cell Biol, 18(12), 7064–7074. https://www.ncbi.nlm.nih.gov/pubmed/9819393

Parra, K. J., Keenan, K. L., & Kane, P. M. (2000). The H subunit (Vma13p) of the yeast V-ATPase inhibits the ATPase activity of cytosolic V1 complexes. J Biol Chem, 275(28), 21761–21767. https://doi.org/10.1074/jbc.M002305200

Paulo, J. A., O’Connell, J. D., Gaun, A., & Gygi, S. P. (2015). Proteome-wide quantitative multiplexed profiling of protein expression: carbon-source dependency in Saccharomyces cerevisiae. Mol Biol Cell, 26(22), 4063–4074. https://doi.org/10.1091/mbc.E15-07-0499

Pettersen, E. F., Goddard, T. D., Huang, C. C., Couch, G. S., Greenblatt, D. M., Meng, E. C., & Ferrin, T. E. (2004). UCSF Chimera--a visualization system for exploratory research and analysis. J Comput Chem, 25(13), 1605–1612. https://doi.org/10.1002/jcc.20084

Roh, S. H., Stam, N. J., Hryc, C. F., Couoh-Cardel, S., Pintilie, G., Chiu, W., & Wilkens, S. (2018). The 3.5-A CryoEM Structure of Nanodisc-Reconstituted Yeast Vacuolar ATPase Vo Proton Channel. Mol Cell, 69(6), 993–1004 e1003. https://doi.org/10.1016/j.molcel.2018.02.006

Sennoune, S. R., Bakunts, K., Martinez, G. M., Chua-Tuan, J. L., Kebir, Y., Attaya, M. N., & Martinez-Zaguilan, R. (2004). Vacuolar H+-ATPase in human breast cancer cells with distinct metastatic potential: distribution and functional activity. Am J Physiol Cell Physiol, 286(6), C1443–1452. https://doi.org/10.1152/ajpcell.00407.2003

Seol, J. H., Shevchenko, A., Shevchenko, A., & Deshaies, R. J. (2001). Skp1 forms multiple protein complexes, including RAVE, a regulator of V-ATPase assembly. Nat Cell Biol, 3(4), 384–391. https://doi.org/10.1038/35070067

Sharma, S., Oot, R. A., & Wilkens, S. (2018). MgATP hydrolysis destabilizes the interaction between subunit H and yeast V1-ATPase, highlighting H’s role in V-ATPase regulation by reversible disassembly. J Biol Chem, 293(27), 10718–10730. https://doi.org/10.1074/jbc.RA118.002951

Smardon, A. M., Diab, H. I., Tarsio, M., Diakov, T. T., Nasab, N. D., West, R. W., & Kane, P. M. (2014). The RAVE complex is an isoform-specific V-ATPase assembly factor in yeast. Mol Biol Cell, 25(3), 356–367. https://doi.org/10.1091/mbc.E13-05-0231

Smardon, A. M., Nasab, N. D., Tarsio, M., Diakov, T. T., & Kane, P. M. (2015). Molecular Interactions and Cellular Itinerary of the Yeast RAVE (Regulator of the H+-ATPase of Vacuolar and Endosomal Membranes) Complex. J Biol Chem, 290(46), 27511–27523. https://doi.org/10.1074/jbc.M115.667634

Smardon, A. M., Tarsio, M., & Kane, P. M. (2002). The RAVE complex is essential for stable assembly of the yeast V-ATPase. J Biol Chem, 277(16), 13831–13839. https://doi.org/10.1074/jbc.M200682200

Smith, A. N., Skaug, J., Choate, K. A., Nayir, A., Bakkaloglu, A., Ozen, S., Hulton, S. A., Sanjad, S. A., Al-Sabban, E. A., Lifton, R. P., Scherer, S. W., & Karet, F. E. (2000). Mutations in ATP6N1B, encoding a new kidney vacuolar proton pump 116-kD subunit, cause recessive distal renal tubular acidosis with preserved hearing. Nat Genet, 26(1), 71–75. https://doi.org/10.1038/79208

Su, Y., Blake-Palmer, K. G., Sorrell, S., Javid, B., Bowers, K., Zhou, A., Chang, S. H., Qamar, S., & Karet, F. E. (2008). Human H+ATPase a4 subunit mutations causing renal tubular acidosis reveal a role for interaction with phosphofructokinase-1. Am J Physiol Renal Physiol, 295(4), F950–958. https://doi.org/10.1152/ajprenal.90258.2008

Sumner, J. P., Dow, J. A., Earley, F. G., Klein, U., Jager, D., & Wieczorek, H. (1995). Regulation of plasma membrane V-ATPase activity by dissociation of peripheral subunits. J Biol Chem, 270(10), 5649–5653. https://doi.org/10.1074/jbc.270.10.5649

Sun-Wada, G. H., Wada, Y., & Futai, M. (2003). Lysosome and lysosome-related organelles responsible for specialized functions in higher organisms, with special emphasis on vacuolar-type proton ATPase. Cell Struct Funct, 28(5), 455–463. https://doi.org/10.1247/csf.28.455

Vasanthakumar, T., Bueler, S. A., Wu, D., Beilsten-Edmands, V., Robinson, C. V., & Rubinstein, J. L. (2019). Structural comparison of the vacuolar and Golgi V-ATPases from Saccharomyces cerevisiae. Proc Natl Acad Sci U S A, 116(15), 7272–7277. https://doi.org/10.1073/pnas.1814818116

Zhao, J., Benlekbir, S., & Rubinstein, J. L. (2015). Electron cryomicroscopy observation of rotational states in a eukaryotic V-ATPase. Nature, 521(7551), 241–245. https://doi.org/10.1038/nature14365

