## Supplemental Figure 1 for "Chimeric a-subunit isoforms generate functional yeast V-ATPases with altered regulatory properties *in vitro* and *in vivo*"

**Supplemental Figure 1:** A. Structures of Vph1NT and Stv1NT indicating residues previously implicated in PI(3,5)P2 binding (pink residues in Vph1NT) and PI(4)P binding (yellow residues in Stv1NT. B. Sequence alignment of Vph1NT and Stv1NT. The proximal domain sequences of both isoforms are highlighted in orange and the distal domain sequences are in green. The PI(4)P site in Stv1NT is boxed and indicated by a yellow circle and the PI(3,5)P2 site in Vph1NT is boxed and indicated by a pink circle. C. Gel filtration traces for the indicated NT constructs after FLAG purification. As described previously, there are dimer and monomer peaks; the monomer fraction was used for all subsequent assays.

Supplemental Figure 1

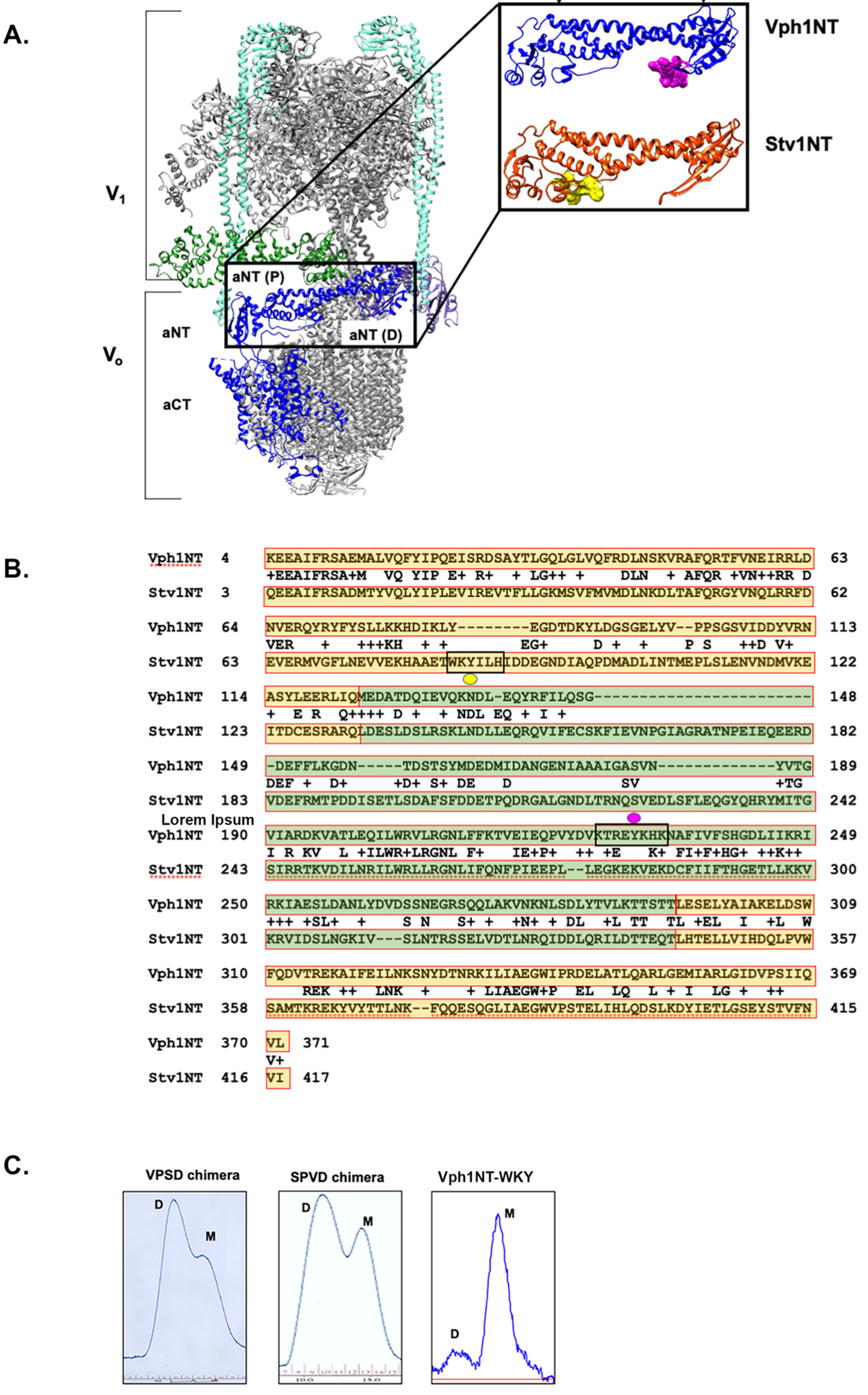
